# Signatures of digital Polycomb regulation in functional iPSC heterogeneity between individuals

**DOI:** 10.1101/2025.07.25.666753

**Authors:** Ander Movilla Miangolarra, Julia Kurlovich, Fereshteh Torabi, Oleksandr Dovgusha, Yang Cao, Lihan Lin, Ufuk Günesdogan, Stefan Schoenfelder, Martin Howard

## Abstract

Induced pluripotent stem cells (iPSCs) show substantial heterogeneity between individuals in their capacity to differentiate into various cell types, hindering their translational application. To explore the causes of this variability, we analysed a panel of ten human iPSC lines, evaluating their differentiation efficiency for two different fates, profiling their transcriptome and chromatin state. Using a machine learning approach, we related chromatin state variability with transcriptional differences. We found that regulation of the Polycomb-mediated histone modification H3K27me3 at specific gene loci exhibited the most consistent differences among iPSC lines, frequently displaying a digital ON/OFF pattern, consistent with a mathematical model based on Polycomb read-write feedback. Notably, altered gene expression patterns of Polycomb-regulated genes were largely propagated into an intermediate state in the differentiation trajectory. Finally, we uncovered a subset of Polycomb-regulated genes with heterogeneous expression and chromatin signatures that were highly predictive of iPSC differentiation efficiencies for two cell types.

## Introduction

All cells in a multicellular organism originate from a totipotent zygote. In mammals, the zygote develops into the blastocyst containing the pluripotent epiblast. As epiblast cells undergo differentiation, they progressively exit the pluripotent state and mature into terminally differentiated cell types. In the past two decades, strategies to induce pluripotency in terminally differentiated cells have attracted much attention [1]. These methods involve the ectopic expression of core pluripotency-associated transcription factors that reprogram somatic cells into induced pluripotent stem cells (iPSCs). iPSCs enable the *in vitro* recapitulation of developmental processes, making them a cornerstone of regenerative cell therapies and disease modelling [2].

However, there is substantial variability in the ability of human iPSC (hiPSC) lines to differentiate into target cell types [3–7]. This heterogeneity has been linked to genetic variability between cell lines and, specifically, between donors [8–10], as well as to differential DNA methylation [3, 5, 11]. Interestingly, genes targeted by the histone-modifying Polycomb complex are prone to considerable expression variability in hiPSCs derived from the same donor [12], suggesting a heterogeneous but significant influence of chromatin regulators. Furthermore, another study showed pronounced variability in histone post-translational modifications (PTMs) across hiPSC lines, exceeding transcriptomic variability [13].

Despite these studies, the extent to which chromatin regulation contributes to the differentiation efficiencies of pluripotent cell lines remains unclear. To better understand the effects of chromatin heterogeneity, we analysed differentiation properties, transcriptome and chromatin features (accessibility and histone modifications) for a panel of ten feeder- and transgene-free, fibroblast-derived, male and female hiPSC lines generated by the HipSci consortium [14].

In this study, we focused on differentially expressed genes (DEGs) across the ten hiPSC lines and sought to relate this differential expression to chromatin accessibility and histone PTM levels. To this end, we developed a computational analysis pipeline based on a support vector machine (SVM). Unlike other machine learning approaches that primarily aim to predict gene expression from epigenomic data (e.g., [15, 16]) or to classify chromatin states (e.g., ChromHMM [17]), our focus was on uncovering the logic of chromatin-mediated transcriptional regulation.

Our analysis shows that the behaviour of around 70% of DEGs could be related to changes in the levels of at least one of the profiled histone marks. We found that the genes whose repression was related to H3K27me3, which is deposited by Polycomb Repressive Complex 2 (PRC2, [18]), displayed the most consistent differences among cell lines. Importantly, we identified a subset of the H3K27me3-regulated DEGs that are predictive of the differentiation efficiencies of the different cell lines into two cell types. Moreover, the regulation of H3K27me3 showed a digital, switch-like behaviour that was consistent with a mathematical model based on read-write feedback of the Polycomb complex [18–21]. Furthermore, we found that the aberrant expression levels of dysregulated Polycomb target genes in hiPSCs were largely propagated into an intermediate state along the differentiation trajectory (mesendoderm precursor cells), pointing to an epigenetic memory system acting during early cell fate choice. Taken together, our results highlight the crucial role of Polycomb complexes in regulating the chromatin landscape of pluripotent stem cells, which in turn profoundly influences their differentiation efficiencies.

## Results

### Marked differences in the developmental efficiency of hiPSC lines

A previous study had highlighted the variability among hiPSC lines in their differentiation efficiency for dopaminergic neurons (DNs, [7]). To test whether these differences were specific to the neuronal lineage, we selected ten hiPSC lines from the HipSci consortium that were reprogrammed from skin fibroblasts of nine healthy donors (five male and four female; Supplementary Table 1). We evaluated their competence to generate primordial germ cell-like cells (PGCLCs), for which they were first induced into mesendoderm precursors (pre-ME) and subsequently re-aggregated into embryoid bodies (EBs) in the presence of cytokines [22] (Fig. 1A). On day 4 of differentiation, EBs were analysed by immunostaining for the PGCLC markers AP2Ɣ, BLIMP1 and SOX17 (Fig. 1B, Supplementary Fig. 1A). This revealed marked differences in PGCLC differentiation efficiency among the cell lines: one cell line (*Podx1*) showed high efficiency (∼45% PGCLCs), five exhibited intermediate efficiency (∼15-28%), and two (*Kucg2* and *Letw5*) had low efficiency (∼5%) (Fig. 1B, C, Supplementary Fig. 1A). Notably, two cell lines (*Sojd3* and *Yoch6*) failed to form EBs after repeated attempts (Fig. 1C, D). We also performed FACS (fluorescence-activated cell sorting) on selected cell lines using antibodies against the PGCLC surface markers INTEGRINɑ6 (ITGA6) and EpCAM [23–25], which independently confirmed the immunofluorescence data (Supplementary Fig. 1B). Together, these results demonstrate significant inter-line variability in the differentiation ability of hiPSCs into PGCLCs.

**Figure 1.**
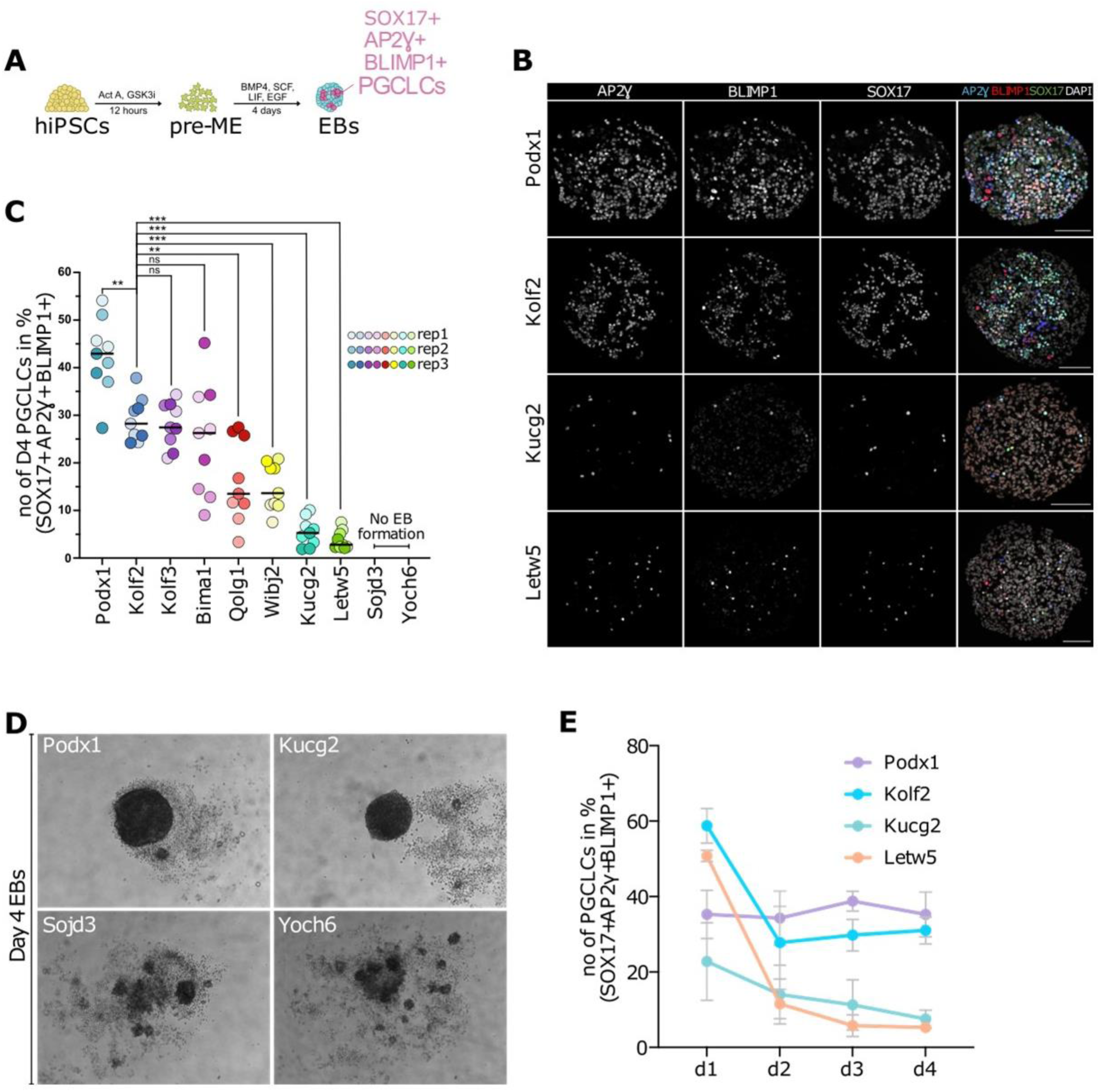
Variation in PGCLC differentiation efficiency. A) Schematic representation of PGCLC differentiation from hiPSCs. B) Immunofluorescence images for AP2Ɣ, BLIMP1 and SOX17 of EBs at day 4 of differentiation from indicated hiPSC lines. AP2Ɣ+, BLIMP1+ and SOX17+ cells represent PGCLCs. C) Quantification of immunofluorescence data showing the percentage of AP2Ɣ+, BLIMP1+ and SOX17+ cells. Median values are represented by horizontal black lines. D) Brightfield images of day 4 (d4) EBs generated from indicated hiPSC lines. E) Quantification of immunofluorescence images showing the percentage of PGCLCs (AP2Ɣ+ BLIMP1+ SOX17+ cells) generated in day (d) 1-4 EBs. Error bars indicate standard deviation, n = 2 biological independent replicates. Scale bars, 100 μm. ActA, Activin A; GSK3i, glycogen synthase kinase 3 inhibitor; BMP4, bone morphogenetic protein 4; SCF, stem cell factor; LIF, leukaemia inhibitory factor; EGF, epidermal growth factor. Two-sided, unpaired t-test: **p <0.01; ***p <0.001; ns, non-significant.

We next asked whether the low number of PGCLCs derived from *Kucg2* or *Letw5* hiPSCs is due to inefficient PGC specification or a failure to maintain the germ cell fate. To address this question, we induced PGCLCs and collected EBs on days one to four for immunofluorescence analysis. EBs derived from *Letw5* hiPSCs initially contained a high proportion of PGCLCs, comparable to cell lines with intermediate or high efficiency (Fig. 1E, Supplementary Fig. 2). However, *Letw5*-derived PGCLC numbers progressively declined over the time course, suggesting a failure to maintain cell identity and/or impaired proliferation. In contrast, EBs derived from *Kucg2* hiPSCs displayed a low number of PGCLCs already at day one of differentiation, indicating inefficient specification of PGCLCs. These findings suggest that *Kucg2* hiPSCs have limited developmental competence to generate PGCLCs, while *Letw5* hiPSCs are capable of PGCLC specification but fail to sustain the germ cell fate, pointing to a defect in fate maintenance rather than in initial developmental efficiency.

To determine whether the impaired differentiation efficiency for PGCLC fate of *Letw5* and *Kucg2* hiPSCs, and the failure of *Sojd3* and *Yoch6* hiPSCs to generate EBs, is specific or reflects a broad impairment of pluripotency, we differentiated six of these hiPSC lines into definitive endoderm (DE). For this, hiPSCs were first induced into mesendoderm, which was then cultured in the presence of Activin A and BMP inhibitor for an additional 48 hours to promote DE differentiation ([22]; Supplementary Fig. 3A). The resulting DE cells were analysed using immunofluorescence for SOX17, which is a marker for both PGCLCs and DE, and the DE marker FOXA2 [26, 27]. Intriguingly, the cell lines that efficiently generated PGCLCs (*Podx1*, *Kolf2*) exhibited relatively low DE differentiation efficiency (∼10-20% SOX17+ and FOXA2+ cells). In contrast, the other cell lines (*Kucg2, Letw5, Sojd3, Yoch6*) differentiated efficiently into DE (40-80% SOX17+ and FOXA2+ cells; Supplementary Fig. 3B, C), which is consistent with a previous study showing high DE differentiation efficiency of *Kucg2* and *Sojd3* hiPSCs [28]. These results were confirmed using FACS (Supplementary Fig. 3D). It is noteworthy that *Kucg2* and *Letw5* hiPSCs robustly upregulated *SOX17* during DE differentiation but failed to do so during PGCLC induction. These results demonstrate that a single hiPSC line can be directed into multiple lineages with variable efficiencies.

### Transcriptional and epigenetic variation among ten hiPSC lines

To further analyse the variation among hiPSC cell lines, we performed RNA-seq (RNA-sequencing). This confirmed a high degree of correlation in gene expression (measured in transcripts per million, TPMs) between cell lines (Pearson correlation, excluding sex chromosomes, >0.96; see Supplementary Fig. 4A), as previously reported [13]. However, Principal Component Analysis (PCA) separated the cell lines into two distinct groups, with three cell lines (*Yoch6*, *Sojd3* and *Kucg2*) clustering on the right side of PC1 (24% of variance, Fig. 2A). Remarkably, these three cell lines displayed low differentiation efficiency for PGCLCs (Fig. 1B, C) and DNs ([7]), but high efficiency for DE (Supplementary Fig. 3B-D).

**Figure 2.**
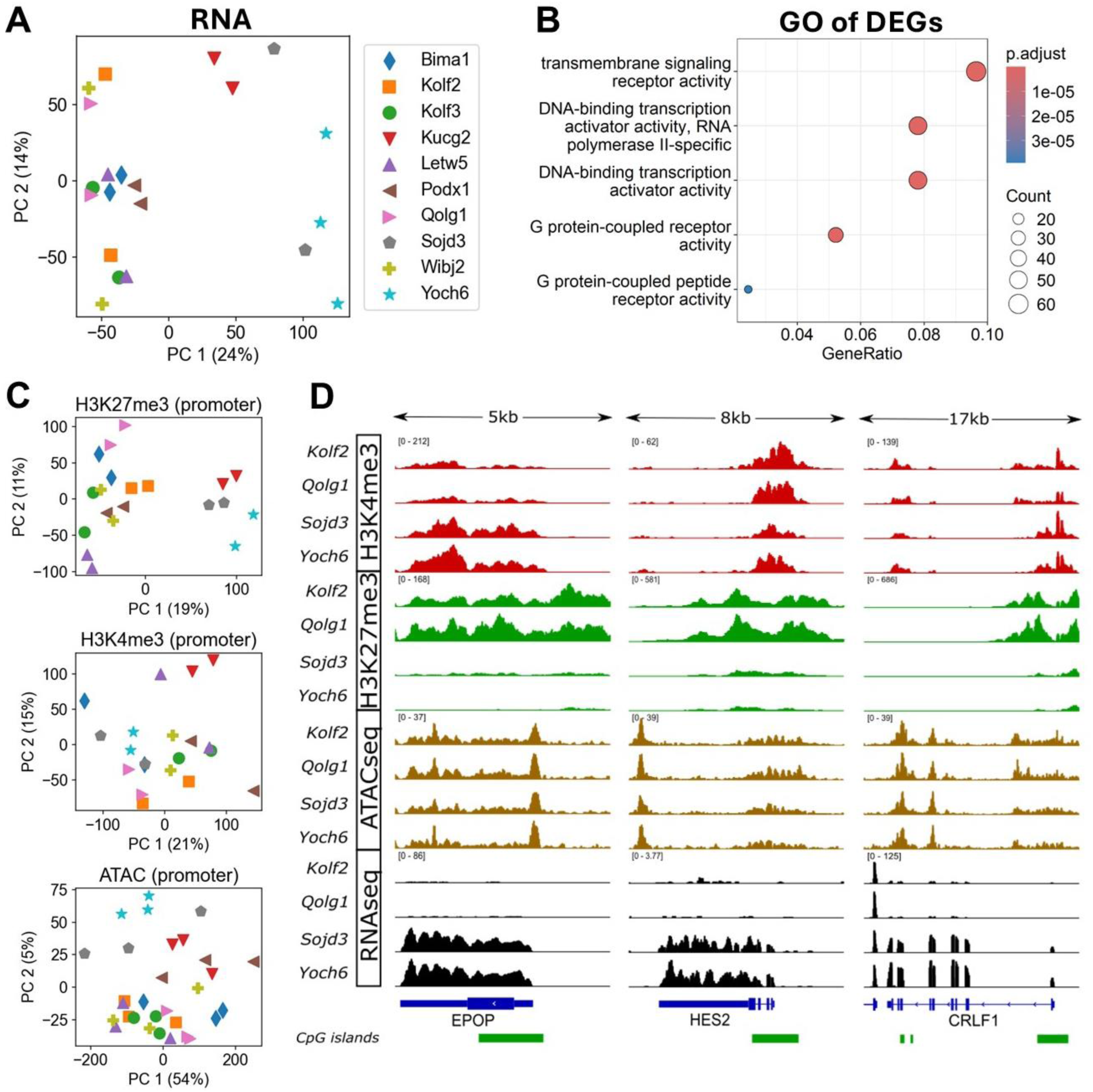
Transcriptional and epigenetic variability in hiPSC lines. A) PCA of the RNA-seq data for the 10 hiPSC lines. Each cell line is represented by a different marker shape and colour (see legend on the right). The variance explained by each of the principal components is given in parenthesis, next to each axis. B) Molecular function GO of the 712 DEGs. The fraction of genes (out of the total of 712) involved in a given molecular function is represented along the x-axis. C) PCA of the epigenomic profiles. For each gene, the signal was integrated over the promoter region. See legend in panel A. D) Genome browser view of three DEGs, for four of the hiPSC lines. RNA-seq, ATAC-seq and two histone marks (H3K4me3 and H3K27me3) are shown. ATAC-seq data were obtained from [30].

Differential gene expression analysis based on pairwise comparisons between cell lines (Methods) uncovered 712 DEGs on autosomes among the ten hiPSC lines. Gene ontology (GO) analysis of the 712 DEGs for molecular function showed enrichment in transcriptional regulators and transmembrane receptors (Fig. 2B), which could be of importance for developmental processes and maintenance of pluripotency. GO for biological processes supports this view by showing enrichment of genes involved in pattern specification and organ development (Supplementary Fig. 4B).

Next, to gain insights into the chromatin states that may underpin differences in gene expression, we performed Cleavage Under Targets and Tagmentation (CUT&Tag, [29]) for three histone PTMs (H3K4me3, H3K9me3 and H3K27me3) in all ten hiPSC lines (two replicates per histone PTM per cell line). In addition, we obtained chromatin accessibility data (as measured by ATAC-seq) for these cell lines from a previous study [30]. For each epigenetic variable, we performed PCA by integrating the enrichment values at each gene over the promoter region (Fig. 2C). We defined the promoter region as between 1kb up- and downstream of the gene start to better capture the peaks of H3K4me3 and ATAC-seq, similarly to other studies [31]. Of the 4 variables considered (ATAC-seq, H3K4me3, H3K9me3 and H3K27me3), only H3K27me3 mirrored the separation between cell lines observed in PCA using RNA-seq data (Fig. 2A, C). Consistently, local differential enrichment of H3K27me3 correlated with transcriptional activity of DEGs including *EPOP*, *HES2* and *CRLF1* (Fig. 2D). When considering gene bodies rather than promoters (defined as from 1kbp downstream of the gene start to the gene end, for consistency with the previous promoter definition), PCA indicated both repressive histone marks (H3K9me3 and H3K27me3) also mirrored the separation seen in the PCA for RNA-seq data (Supplementary Fig. 4C). However, for H3K9me3, this was primarily along PC2, suggesting a less pronounced segregation. We confirmed that our conclusions are robust to an alternative promoter definition (-1kb to gene start).

Overall, our analyses revealed a substantial number of DEGs between human iPSCs, and showed a genome-wide relation between transcriptional heterogeneity and the levels of repressive histone marks, particularly H3K27me3.

### Linking variation in chromatin features with transcriptional output using SVMs

We next sought to elucidate how variability in histone PTMs or chromatin accessibility relates to transcriptional activity. To address this, we developed a computational pipeline whose main objective was to understand the logic linking chromatin variability to transcriptional regulation.

The pipeline takes as input the values for histone PTM enrichment and chromatin accessibility (hereafter referred to as ‘epigenomic variables’), and the transcription levels for each DEG in each cell line. To avoid artificially inflating the number of datapoints by combining replicates of different epigenomic variables, we used the merged data for each variable in each cell line. Since H3K4me3 and chromatin accessibility typically display peaks near transcription start sites, we quantified their signals over promoter regions (see Methods for details). The signals for the repressive marks H3K9me3 and H3K27me3 were integrated across the entire gene (from start to end) [32]. We also tried using alternative genomic regions for signal quantification, but the performance of the pipeline remained similar or was slightly reduced (Supplementary Fig. 4D). The cornerstone of the pipeline is a Support Vector Machine (SVM), a machine learning method designed to predict a binary outcome (here, transcription) based on an input (epigenomic variables). A key advantage of this technique is that it is straightforward to interpret the parameters learnt from the data. Since a SVM is a binary classifier, we binarized the transcription values to distinguish between genes with higher or lower expression in each cell line, for which we used a K-means clustering approach (Fig. 3A). After binarization, the data were ready to be passed to the SVM. Due to the differences in gene-specific regulation, we analysed each gene independently, by training a separate SVM for each one. While reducing transcription levels to a binary output may be an oversimplification, this approach allows for robust predictions using relatively sparse datasets, as in this case with ten datapoints (cell lines) per gene.

**Figure 3.**
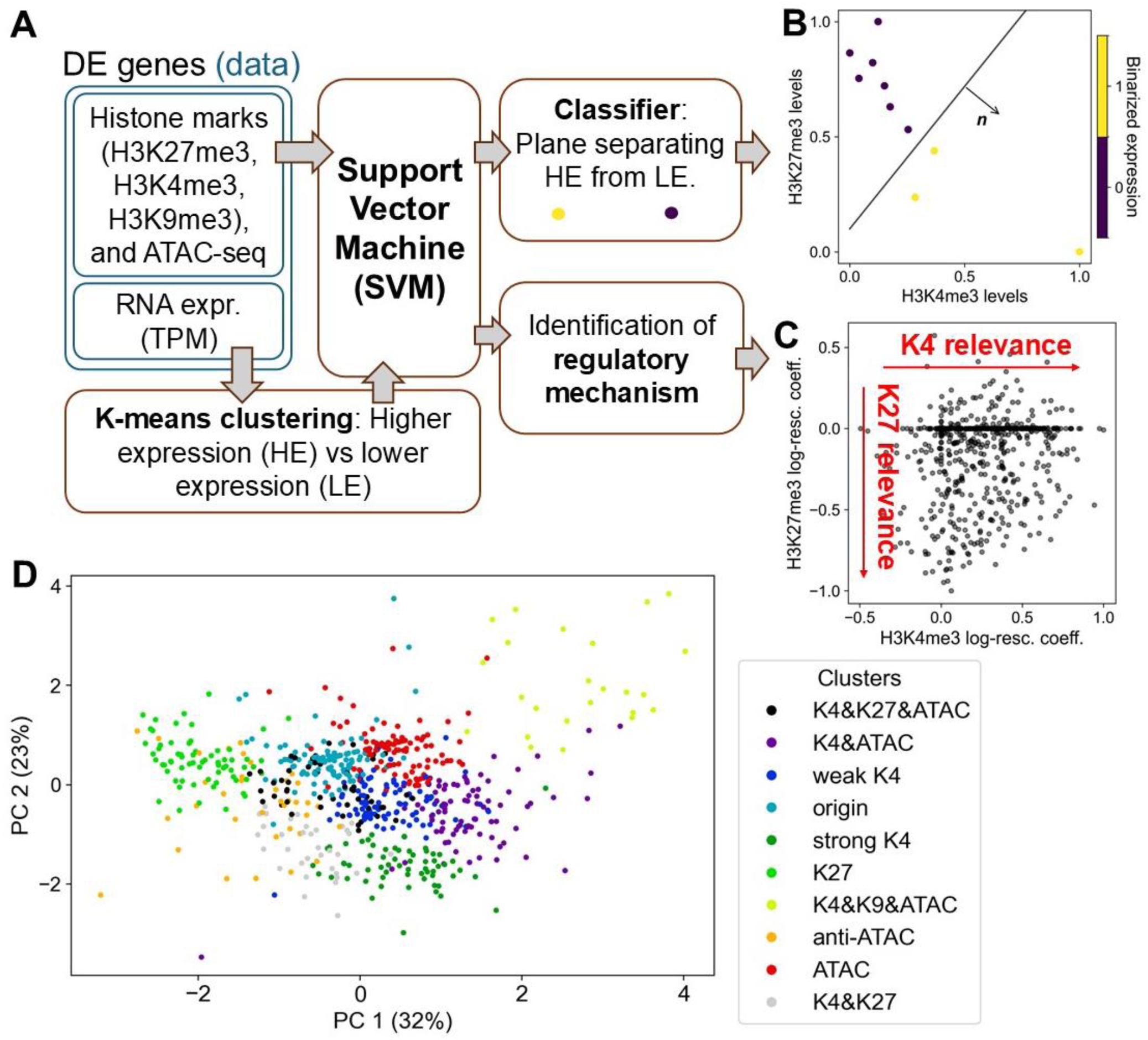
Using an SVM to identify regulatory modes. A) Schematic of the computational pipeline used to analyse the combined datasets. B) Example of a 2D SVM line separating cell lines with higher expression from those with lower expression for a given gene. C) Example of the regulatory space obtained from a 2D SVM. Each point is a gene, whose position in the regulatory space is determined by the relationship between transcription and the epigenomic variables. D) Two-dimensional PC projection of the 4D regulatory space, with the regulatory modes (clusters) colour coded. The variance explained by each of the principal components is given in parenthesis, next to each axis.

Our SVMs work on the space of the input data for each gene (four axes, three for histone PTMs and one for chromatin accessibility). Each cell line is represented as a datapoint within this four-dimensional epigenomic space, alongside the transcriptional output with respect to a given gene. Once the SVM has been trained, the result is an optimal hyperplane that, if possible, separates cell lines with high expression of a gene of interest from those with low expression (for details, see Methods). This hyperplane can then be used to predict the binary expression levels of additional datapoints (or cell lines; Fig. 3B). The vector normal to this hyperplane indicates the direction in which the epigenomic variables must change to upregulate transcription. This vector is our gateway to identifying the regulatory mechanism, mathematically described by ***n***_*i*_ = (*n*_*i*,*K*4_, *n*_*i*,*K*9_, *n*_*i*,*K*27_, *n*_*i*,*ATAC*_). Thus, each gene *i* can be mapped to a point within a regulatory space, which is defined by the extent to which each epigenomic variable contributes to the prediction of transcription, in terms of the log-rescaled ***n***_*i*_ coefficients (see Methods and Fig. 3C). Next, we clustered genes according to their regulatory mode using another K-means algorithm (see Methods and Supplementary Fig. 5 A, B), resulting in 10 clusters (named after the variables with average |log − resc. coeff. | > 0.2). About 80% of the DEGs could be clustered into a biologically meaningful regulatory mode involving the chromatin state, with only a minority of about 20% of DEGs in the ‘origin’ cluster, for which we could not establish a strong connection between transcription and epigenomic variables (see Fig. 3D for a two-dimensional PC projection of the resulting four-dimensional regulatory space).

The regulatory space is shown in terms of log-rescaled SVM coefficients, which integrate the SVM output (hyperplane coefficients, ***n***_*i*_) and the variation of the corresponding epigenetic variable at the gene of interest (hence, the ‘rescaled’ prefix). Since the data has to be normalised for robust SVM training, this rescaling ensures that the original scale of variations in histone marks is taken into account. The absolute value of the log-rescaled coefficient is bounded by 1, with values closer to 1 indicating stronger regulatory association. The sign of the coefficient indicates the relationship with transcription: negative values indicate repression, and positive values indicate activation. As expected, the majority of the log-rescaled H3K4me3 coefficients were positive, whereas the majority of the H3K27me3 coefficients were negative, consistent with their respective roles as activating or repressive chromatin marks [33].

Classification of genes by chromatin regulation using a SVM is only meaningful if the SVM itself performs well; that is, if it can correctly classify high or low levels of transcription from its epigenomic signals. To test this, we trained the SVM on data from nine hiPSC lines and tried to predict expression patterns for the tenth independent hiPSC line based on its epigenomic profile. On average, the SVM (and, by extension, the entire pipeline) accurately predicted gene expression 78.4% of the time (95% Confidence Interval [CI]: 63%-88%, Supplementary Fig. 5C; compare with the mean accuracy of a logistic regression approach: 76.9%). Using Platt scaling [34] to obtain a probability associated with each prediction, we confirmed the near-perfect calibration of the SVM (Supplementary Fig. 5D). To rigorously test the accuracy of the classification, we drew a receiver operating characteristic (ROC) curve for each of the cell lines (see

Methods and Supplementary Fig. 5E). When predicting gene expression in the tenth cell line using the remaining nine for training, we obtained an average area under the ROC (AUROC) of 0.832 (95% CI: 0.64-0.95). Since a value of 1.0 represents a perfect classifier and 0.5 a random one, this result suggests that the SVM performs very well, especially given the limited training data available for each gene (nine datapoints). As an alternative approach, we also used a logistic regression instead of an SVM, for which we obtained an average AUROC of 0.828 (95% CI: 0.48-0.96, Supplementary Fig. 5F). When further interrogating this small difference, we noticed that the prediction is harder to make correctly for the three hiPSC lines that separate from the remaining cell lines in the PCA analysis for RNA-seq data (Fig. 2A; Supplementary Fig. 5E, F). In these cell lines, the AUROC of the logistic regression drops as low as 0.6, whereas the SVM still maintains a satisfactory performance of about 0.7. These results demonstrate that the SVM predicts high or low gene expression from epigenomic variables more robustly than a logistic regression, particularly in cases with limited data, with marked improvement in the lower bound of the 95% CI.

### Chromatin accessibility does not always correlate with transcription

We next investigated the differences in the behaviour of genes associated with each regulatory mode in more depth. Our clustering identified 10 distinct regulatory modes. Unexpectedly, chromatin accessibility emerged as a salient feature only in a subset of these modes. Thus, we interrogated the data to clarify the relation between accessibility and transcription in each of the regulatory modes. The average log-rescaled ATAC-seq coefficient for each regulatory mode showed that, while half of the clusters exhibit a positive log-rescaled coefficient, four clusters have coefficients close to zero, indicating minimal or no association between accessibility and gene expression (Fig. 4A). Surprisingly, in one cluster, the ATAC-seq signal at the promoter seemed to correlate with gene repression (named ‘anti-ATAC’). In addition, many genes silenced by H3K27me3 either did not show significant changes in chromatin accessibility, or the changes in accessibility were not correlated with transcriptional variability. In fact, less than 40% of genes repressed by H3K27me3 displayed reduced ATAC-seq signal at their promoters. These results are consistent with reports showing that chromatin accessibility, as measured by ATAC-seq, is not always tightly linked to transcriptional activity [35–37].

**Figure 4.**
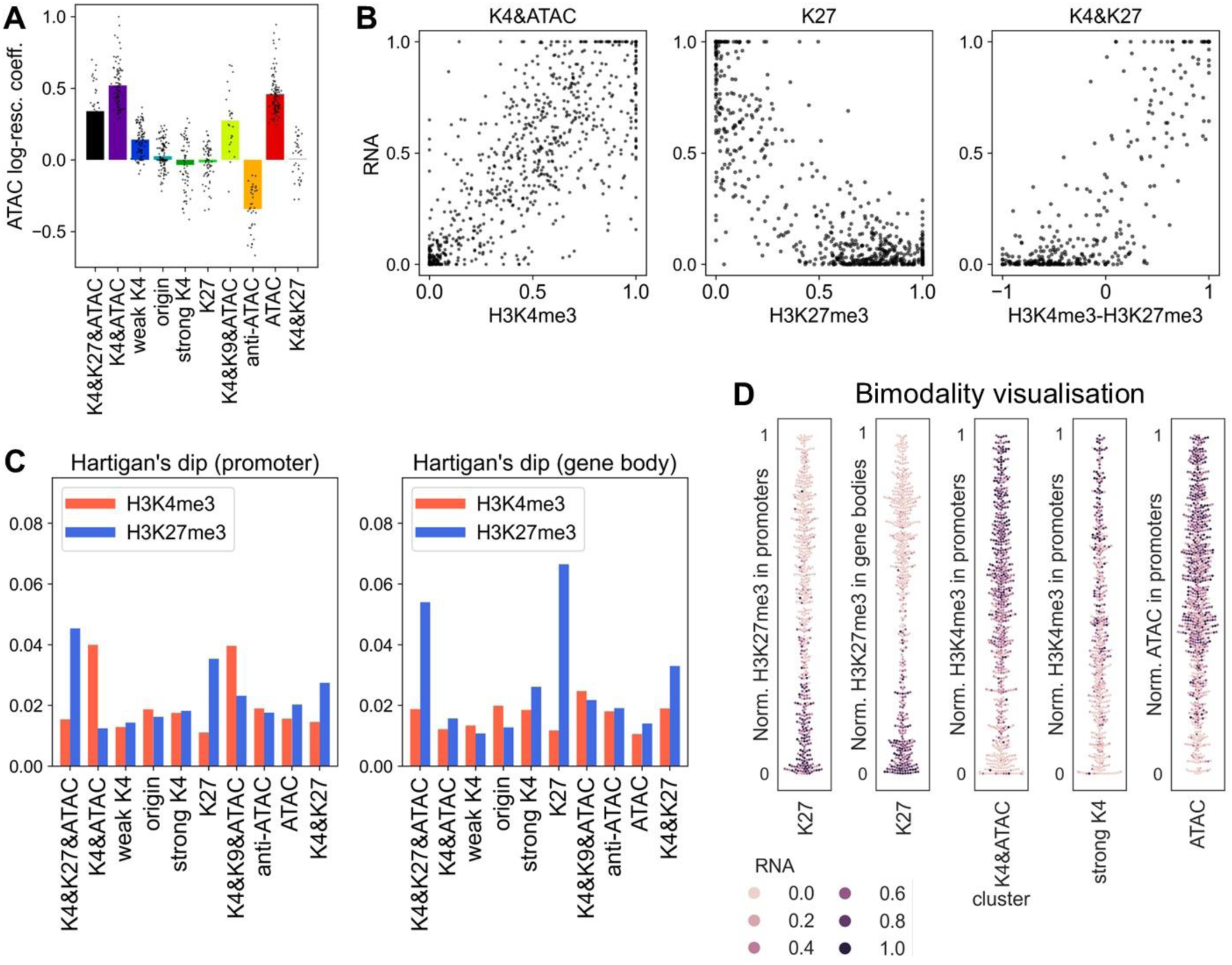
Analysis of regulatory modes. A) Log-rescaled coefficient for accessibility in each of the regulatory modes. Bars indicate averages and dots individual genes. B) Scatter plot of transcriptional output as a function of the relevant epigenomic variables for each regulatory mode. Both transcription and the epigenomic variables are normalised between 0 and 1 (per gene, as explained in the text). C) HDS for the histone PTMs H3K4me3 and H3K27me3, for each regulatory mode, taken over two different annotations: promoter (left panel) and gene body (right panel). For each regulatory mode the H3K4me3 HDS is depicted in red and the H3K27me3 HDS is shown in blue. D) Visualisation of bimodality and its relationship with transcription, for selected regulatory modes. The normalised value of the relevant epigenomic variable of a datapoint is given by its position on the y-axis. The width of the swarm of points shows the number of datapoints in that range of enrichment values, revealing clear bimodality in some cases. The colour of the datapoints represents normalised transcriptional output for a given gene and cell line.

### H3K27me3 regulation is digital

To visualise the data within the different regulatory modes, we combined all genes within each cluster by normalising the values of transcription and the epigenomic variables between 0 and 1 (the maximum of each variable across cell lines was mapped to 1 and the minimum to 0), independently for each gene. Then, all genes within a cluster (each gene with ten datapoints, one from each cell line) were visualised within a single graph (Fig. 4B). We observed a variety of forms of transcriptional regulation: from a fairly linear relationship between RNA expression and H3K4me3 for cluster ‘K4&ATAC’, to a more complex link between RNA expression and two histone marks (‘K4&K27’ cluster). Notably, the ‘K27’ cluster exhibited bimodality, with large numbers of datapoints in the expressed region of the graph (low H3K27me3 and high expression) and in the silent region of the graph (high H3K27me3 and low expression), but with a largely unpopulated central region (Fig. 4B).

Based on these observations, we sought to dissect the nature of the relationship between epigenomic variables and transcription. Do these features vary continuously, or are there abrupt changes? A more rigorous quantitative version of the previous qualitative analysis involves computing Hartigan’s Dip Statistic (HDS), which, given the distribution of a variable (e.g., histone PTM level within a regulatory mode), quantifies the distance to the nearest unimodal distribution [38]. Thus, a higher HDS indicates a greater deviation from unimodality, suggesting that the distribution is bimodal or multimodal and may therefore reflect two or more discrete states. We computed the HDS for H3K4me3 and H3K27me3 using the promoter and the gene body annotation (Fig. 4C, see also Supplementary Fig. 6A for a similar analysis with RNA-seq and ATAC-seq data, which show moderate or no bimodality at all). Overall, we found that H3K27me3 exhibits the highest degree of bimodality, particularly at gene bodies, across all three regulatory modes in which it is involved (see also Supplementary Fig. 6B for the histograms of some of these distributions, with various degrees of bimodality). However, we also found significant bimodality for H3K4me3 in the promoter regions of the ‘K4&ATAC’ and ‘K4&K9&ATAC’ regulatory modes, although the latter consists of only ∼20 genes. Using an alternative promoter definition (from -1kb to gene start), we confirmed that H3K27me3 remains the most bimodal mark and that its bimodality is also greater at the gene body than at the promoter.

We further visualised this bimodality using a “swarmplot” in which the normalised enrichment of the histone mark (for each datapoint of a given regulatory mode) is represented on the y-axis, and the colour denotes the normalised expression (Fig. 4D). Interestingly, in the case of H3K27me3 at gene bodies, within the ‘K27’ regulatory mode, bimodality seems to be closely linked to transcriptional activity: the top lobe of the distribution corresponds to repressed instances, while the lower lobe represents expressed ones. A sharp transition in expression occurs around 30% of the maximum H3K27me3 signal, marked by a narrow intermediate region containing relatively few datapoints. The fact that the transition from repressed to expressed occurs in a region in between the two main lobes of H3K27me3 enrichment (as opposed to taking place in the middle of one of the lobes) suggests a role for H3K27me3 bimodality in regulating transcription. Furthermore, the transition value of 30% enables repression to be stably maintained even after DNA replication, when, on average, histone modification levels will be transiently halved [39]. It should also be noted that other epigenomic variables do not exhibit pronounced bimodal behaviour (see two examples in Fig. 4D, ‘strong K4’ and ‘ATAC’). Even in cases where H3K4me3 shows some degree of bimodality (e.g. ‘K4&ATAC’ cluster), the bimodality is less marked and the transition from repressed to expressed states appears more gradual.

### The behaviour of H3K27me3 can be explained using a simple mathematical model

Above, we have highlighted the bimodal behaviour of H3K27me3 in gene bodies and its tight relationship with transcriptional output. However, we also observed a crosstalk between H3K27me3 and H3K4me3 in at least two regulatory modes, with similar but slightly different behaviour in terms of multimodality or correlations between the marks. In addition, we found that the behaviour of H3K27me3 in promoter regions may be less bimodal than in gene bodies. We therefore attempted to explain these results mechanistically using a mathematical model.

The model considers each gene to be an independent locus where histone 3 tails at promoters can be in one of five possible states: H3K4me3, unmodified, H3K27me1, H3K27me2 and H3K27me3. At gene bodies, histone 3 tails can only be methylated at residue 27 (for a schematic of the model see Fig. 5A). Transition between states take place via basal linear rates (parametrised by *k*_0_) or, for the addition of a methyl group to the lysine 27, via the allosteric feedback of PRC2, resulting in a nonlinear transition rate proportional to the fraction of H3K27me3 in the locus, parametrised by *k*_*f*_. So far, this largely follows other well-established mathematical models [20, 21, 40, 41], except for the fact that we considered promoter regions and gene bodies separately. These two sets of histone tails only interact via the Polycomb feedback, controlled by the parameter *c* (potentially reflecting physical interactions between distal sites within the gene). We take *k*_*f*_ equal for promoters and gene bodies. However, we assume that promoter regions have a larger basal rate due to their greater overlap with CpG islands, which are known to recruit Polycomb complexes. To reflect this, we set *k*_0,*p*_ = 1.5*k*_0,*b*_, where *k*_0,*p*/*b*_ is the basal rate of promoters/gene bodies. In addition, we also considered a periodic perturbation due to DNA replication, where all histone PTMs are diluted by an average factor of two due to the inclusion of newly synthesised unmethylated histones. This has been shown to be important for transcriptional regulation, both experimentally [42–44] and theoretically [45, 46]). For more details on the mathematical modelling, see Methods.

**Figure 5.**
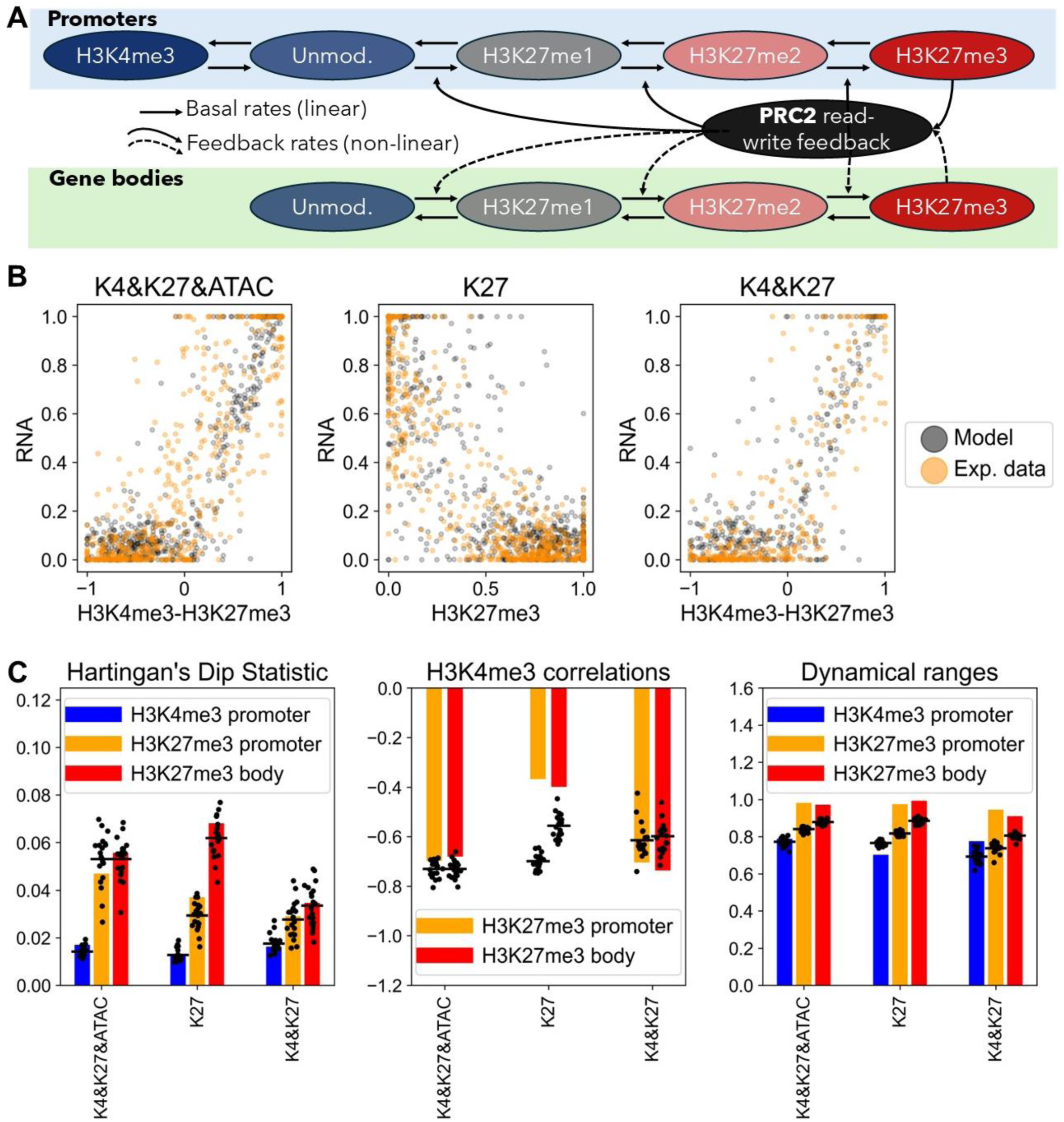
A mathematical model recapitulates digital H3K27me3 states. A) Schematic of the mathematical model. B) Qualitative comparison between synthetic data (grey) and experimental data (orange), for the three regulatory modes involving H3K27me3. C) Quantitative comparison between model and experimental data for the same regulatory modes as in B). From left to right: The first panel depicts the HDS in experimental data (bars) and the model (black points, one for each of the 20 stochastic simulations, and black line for the mean of simulations). HDS is shown for three different combinations of histone mark and region of the gene. Middle: Correlation between H3K4me3 at promoters and H3K27me3 at promoters or gene bodies, for three different regulatory modes. Right: Dynamical ranges for each of the histone marks, taken as a fraction of the difference between the maximum and the minimum value of a variable (among the ten cell lines), divided by the maximum.

To compare the output of the model to the experimental data, we used an approach that creates synthetic data for qualitative and quantitative comparison. Furthermore, we assumed that most of the noise in the analysis of experimental data came from 1) technical/biological noise and 2) grouping genes (which may have different features) together, as is done when clustering into regulatory modes. Therefore, we opted for deterministic modelling using ordinary differential equations, to which noise is added before (in the parameters, representing slightly different characteristics of genes) and after (in the resulting outputs, representing technical/biological noise in experiments). Finally, the model required a variable parameter to explain the differences between cell lines. In this case, we chose *E*, which controls the rates at which methyl groups are added to H3K27 residues. This was chosen because the largest cell-line dependent effects occurred in H3K27me3-related regulatory modes (see next section). To obtain data for a given regulatory mode, we simulated the model the same number of times as there are genes clustered into that regulatory mode. Each simulation used different random parameters with a prescribed mean (see Table 1). To simulate each cell line, we modulated the parameter *E* from 1 to 0.8 in ten equally spaced intervals (equal to the number of cell lines). By gathering data from each simulation and adding noise to the outputs, we obtained as many synthetic datapoints as there were experimental ones, which we then analysed in the same manner (Fig. 5B; for more details, see Methods).

**Table 1.** Parameters of the mathematical model, for each of the regulatory modes modelled. In boldface, the parameters that change depending on the regulatory mode.

| | $k_{0,p}$ ($\text{hr}^{-1}$) | $k_f$ ($\text{hr}^{-1}$) | $c$ | $k_{da}$ ($\text{hr}^{-1}$) | $k_{dm}$ ($\text{hr}^{-1}$) | $k_a$ ($\text{hr}^{-1}$) | $T$ (hr) |
| --- | --- | --- | --- | --- | --- | --- | --- |
| K4&K27&ATAC | 0.42 | 3.6 | 0.3 | 2 | 1 | 0.8 | 22 |
| K27 | 0.42 | 3.6 | 0.03 | 2 | 1 | 0.8 | 22 |
| K4&K27 | 0.52 | 3.2 | 0.3 | 2 | 1 | 0.8 | 22 |

With appropriate parameterisation, we found that the model could largely explain the behaviour of the three relevant regulatory modes by only varying two parameters: *c* and *R* (the ratio between the feedback and basal rates *k*_*f*_/*k*_0,*p*_). The agreement was both qualitative (Fig. 5B) and quantitative (Fig. 5C).

Indeed, the model could quantitatively recapitulate the HDS for three different variables including H3K4me3 at promoters, H3K27me3 at promoters and H3K27me3 in gene bodies, and in the three H3K27me3 regulatory modes (Fig. 5C, left panel). In general, gene bodies had the highest HDS (both in experiments and in the model) which, in the model, was due to having a higher *R*. The difference between the HDS in gene bodies and promoters was maximal in the ‘K27’ cluster, which we parametrised as having a lower *c*. Thus, this decoupling of promoter and gene body explains the large difference in behaviour. Increasing *c* makes the HDS of promoters and gene bodies converge, as in the ‘K4&K27&ATAC’ cluster. Finally, it is worth noting that the H3K4me3 HDS was very low in both experiments and in the model. This was surprising, since the two histone PTMs cannot coexist in the same histone tail in the model. Therefore, H3K4me3 levels should respond accordingly to changes in H3K27me3 and reflect its bimodality. However, since H3K4me3 levels respond to changes in all H3K27me PTMs, the resulting bimodality is substantially weaker (H3K27me1 is not as bimodal as H3K27me3) and the added noise further reduces this residual bimodality.

Fig. 5C shows the dynamic ranges of these variables in the model and the experiments, which show generally good agreement. Fig. 5C also demonstrates the correlation between H3K4me3 at promoters and H3K27me3 at gene bodies or promoters for each of the three clusters considered. This correlation was heavily constrained by the exclusive nature of the H3K4 and H3K27 marks, which correctly predicted the quantitative value of the anticorrelation in the ‘K4&K27&ATAC’ and ‘K4&K27’ clusters, but overestimated that of the ‘K27’ cluster, especially at promoters. This is probably due to spatial inhomogeneities that we neglected beyond a separation into promoter and gene body. Nevertheless, the overall quantitative agreement between model and experimental data is very good, especially considering the relatively few parameters in the model and the varied set of quantitative outputs it explains.

Taken together, the model highlights the critical role of the feedback rate in generating switch-like behaviour in histone PTM regulation. Promoters therefore exhibited a lower HDS than gene bodies, with the extent of this difference depending on the coupling *c* between promoter and gene body. Crucially, the model was capable of creating synthetic data, quantitatively and qualitatively similar to the experimental data, by assuming a modest modulation of *E*, the parameter that controls the overall H3K27me levels. Indeed, one of the most important consequences of the switch-like regulation of histone PTMs is that even a modest variation in a parameter can cause a substantial change in histone PTM levels.

### A major role for H3K27me3 in hiPSC transcriptional heterogeneity

Given the marked transcriptional variations observed among cell lines (Fig. 2), we next asked if these differences were uniformly distributed across regulatory modes or whether they could be attributed to just a subset of them. To address this, we obtained heatmaps of the RNA expression (z-score, to standardise the mean and variance of each gene independently) of all ten cell lines for all genes in each cluster, using hierarchical clustering to organise genes and cell lines (Fig. 6A and Supplementary Fig. 7A). The ‘K27’ cluster shows distinctive behaviour with a very clear difference between *Kucg2*, *Sojd3* and *Yoch6* and the remaining hiPSC lines, which have opposite expression patterns in most genes (Fig. 6A). Comparable behaviour for these three cell lines is seen for the other two clusters involving H3K27me3 (Supplementary Fig. 7A). Although a similar trend was also apparent in the ‘K4&ATAC’ cluster, it was substantially less pronounced than in H3K27me3-associated clusters. We calculated the absolute value of the pairwise Pearson correlation of gene expression between two cell lines to assess which regulatory modes show more robust transcriptional differences. Higher values indicate stronger correlation or anticorrelation and, thus, stronger differences between cell lines. This analysis confirmed that the three clusters involving H3K27me3 show the strongest patterns (Fig. 6B), suggesting that this histone PTM plays a key role in the regulation of transcriptional variability in hiPSC lines. In Fig. 2D, we already saw a similar genome-wide relation for H3K27me3; here, we dissected the genes that contribute the most to this relation (see Supplementary File 1 for a list of DEGs per regulatory mode).

**Figure 6.**
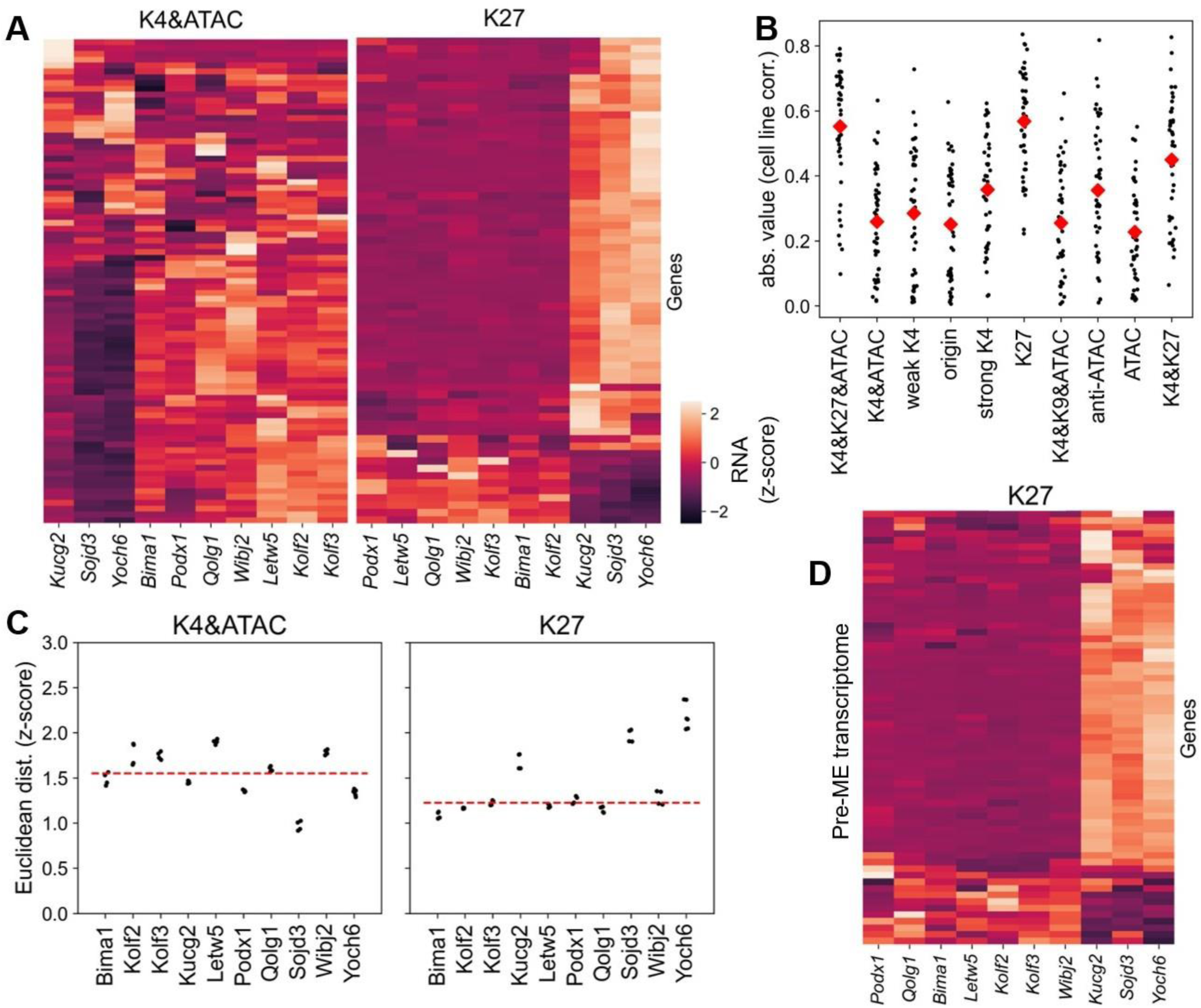
Heterogeneity within regulatory modes between cell lines. A) Heatmap of the transcriptional output (z-score), with purple/black representing repressed genes for a given cell line, and light orange colours representing expressed ones. B) Quantification of cell-line dependent effects, in terms of the absolute value of the pairwise correlations between cell lines, considering only genes within a given regulatory mode. Within each regulatory mode (x axis), each pairwise correlation is depicted by a black dot and the average across all 45 possible pairwise correlations is given by the red diamond. C) Euclidean distance, in terms of the z-scores, between the hiPSC lines and H9 hESCs, using only the genes from a given regulatory mode (specified in the title of each plot). The red dashed line denotes the median distance from hiPSC lines to the hESC line. D) Transcriptional output (normalised to z-scores) for genes in the K27 cluster for the 10 cell lines in the pre-ME state. Same as A) but using the expression values of the cell lines in pre-ME state. Colour bar shown in panel A).

In addition, we compared the transcriptomes of the hiPSC panel with that of a reference human embryonic stem cell (hESC) line (H9 line, from the ENCODE database [47]). The Euclidean distance calculated on z-scored expression values did not show significant differences in the distance between the hiPSC lines and the hESCs when taking all DEGs into account (Supplementary Fig. 7B). However, when dividing the DEGs into the different regulatory modes, striking differences appeared (Fig. 6C). In particular, the ‘K27’ cluster showed a very marked pattern that mirrored the genome-wide transcriptional heterogeneity shown in Fig. 2A (for completeness, every cluster is shown in Supplementary Fig. 7C). Furthermore, the only clusters where *Yoch6, Sojd3* and *Kucg2* were the three most distant lines from hESCs were those involving H3K27me3, suggesting aberrant Polycomb regulation in these cell lines. The fact that the cell lines with diverging H3K27me3 regulation had also low (or null) efficiency to give rise to PGCLCs is unlikely to be a coincidence (*p*<0.04, see Methods). By contrast, the fourth cell line with low PGCLC differentiation efficiency (*Letw5*) initially succeeded in inducing PGCLCs but then failed to maintain PGC fate. This suggests that, in this case, a distinct mechanism, independent of H3K27me3 dysregulation, may result in impaired germ cell development.

Overall, these findings suggest that the chromatin mark H3K27me3 plays a major role in transcriptional heterogeneity between hiPSC lines. Moreover, genes regulated by H3K27me3 exhibited particularly clear differences between *Kucg2*, *Sojd3* and *Yoch6,* and the remaining hiPSC cell lines. Crucially, we found that, by comparison with a hESC reference line, their H3K27me3 regulation was aberrant.

### Pre-ME data suggest inheritance along the developmental trajectory

Given the differences in the transcriptome and epigenome identified in this study, and their correlation with differentiation properties, we hypothesised that the heterogeneities present in the iPSC state are propagated along the differentiation trajectory. To test this hypothesis, we sequenced RNA of all ten hiPSC lines after differentiation into pre-ME, which represents an intermediate state during PGCLC or DE differentiation (see Fig. 1A). We found that 60% of the DEGs in pre-ME were also DEGs in hiPSCs (see Supplementary Fig. 8A). Importantly, the clear differences in transcription of PcG targets were maintained in pre-ME (see Fig. 6D for the expression of genes in the K27 cluster, with a pattern very similar to the iPSC results in Fig. 6A). In addition, the gene clusters involving H3K27me3 from iPSCs still present the most robust transcriptional differences among the cell lines in pre-ME (Supplementary Fig. 8B). Finally, histone marks in the iPSC state were also predictive of expression in the pre-ME state, albeit with slightly lower accuracy than for the iPSC state (Supplementary Fig. 8C, D), which may indicate the existence of an epigenetic memory system that is maintained during differentiation.

### A subset of Polycomb targets is predictive of differentiation properties

Given the robust heterogeneities in the clusters involving H3K27me3 and how they *correlate* with PGCLC differentiation, a key next step is to examine whether the transcriptional output of these targets can be used to *predict* differentiation. To answer this question, we built a logistic regression model to predict PGCLC differentiation from the RNA expression values of the 152 targets in the clusters involving H3K27me3. To minimise the risk of overfitting, only reliable targets were retained for a final model (resulting in a total of 109 genes used, see Methods). As expected from our analysis, the model was successful at predicting the ability to differentiate into PGCLCs (91% accuracy [95% CI: 78%-100%] and 0.95 AUROC [95% CI: 0.86-1.0], with an approximately 80/20 data split). However, the PGCLC dataset is limited for training this type of statistical model (there are five times more parameters than datapoints); therefore, we next asked how well the model would generalise.

We used a large-scale study of neuronal differentiation into DNs [7] to address this question. Remarkably, our model trained on our separate PGCLC data and using only 109 genes obtained similar metrics for predicting DN differentiation ability to the model of Ref. [7], which was trained on DN differentiation assays and used the expression levels of a 130-fold greater number of genes, about 13,000 (Fig. 7A). When our model was trained on DN differentiation data, the set of highly predictive PcG targets was narrowed further (only 47 genes, see Supplementary File 2 for their identity and coefficients), accuracy increased (82% [95% CI: 75%-86%]) and the performance metrics were improved substantially (Fig. 7A). Adding characteristics of the cell line, such as age or sex of the donor, did not further improve the performance of the model. Taken together, these results suggest the following: *1)* Using a small number of well-chosen targets (Polycomb-controlled genes in this case) can outperform the use of entire transcriptomes in the prediction of differentiation and *2)* PGCLC and DN differentiation share many of their predictors at the iPSC stage. This is also apparent from the high correlation (*r*=0.71) between the expression of individual genes and differentiation into the two different fates (Fig. 7B).

**Figure 7.**
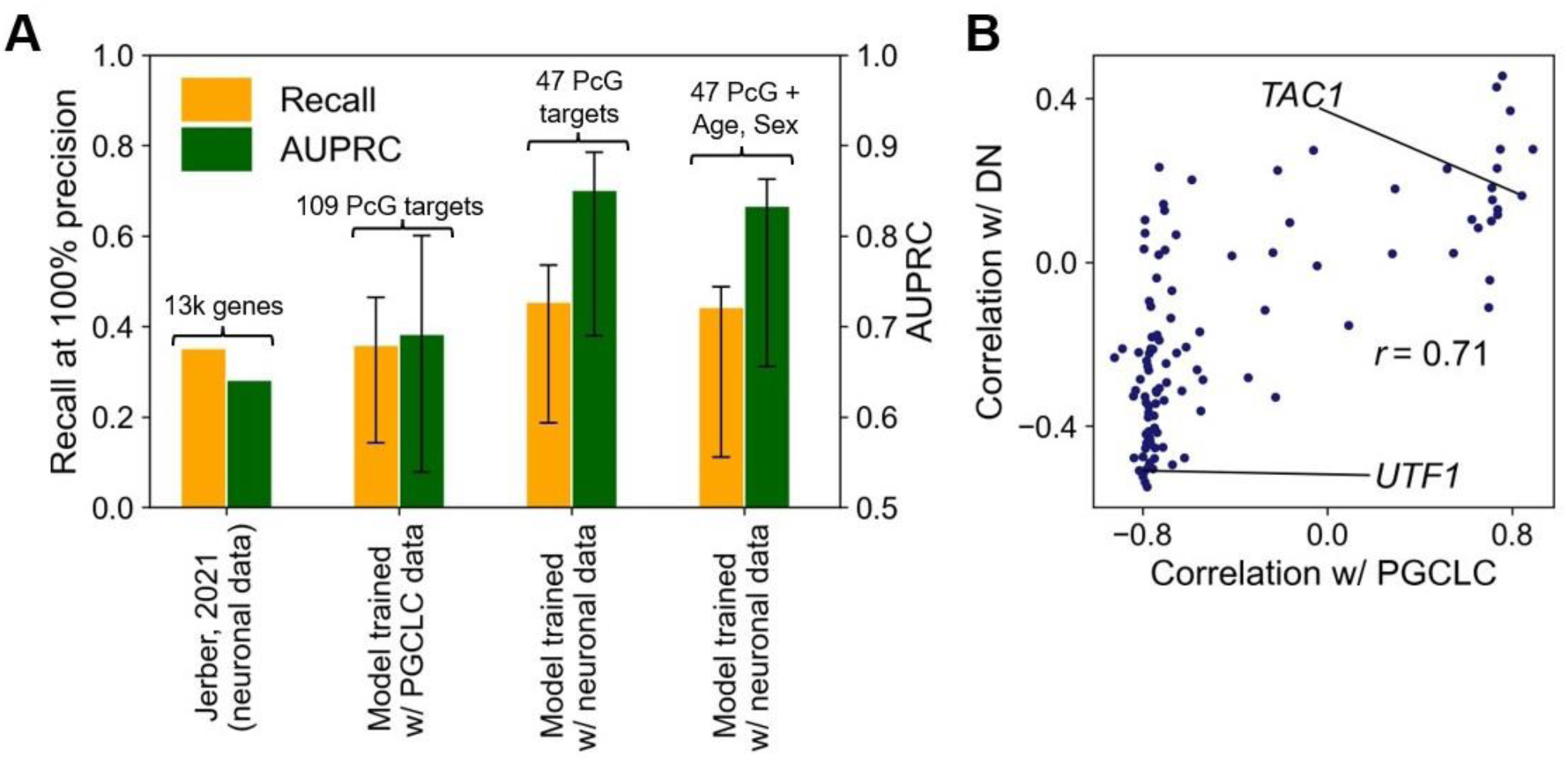
Highly predictive model of differentiation efficiency from PcG target expression. A) Performance metrics for the logistic regression models (using leave-one-out cross-validation), when tested on neuronal differentiation assays (AUPRC, Area Under the Precision-Recall Curve). Both metrics are bounded by 1 and higher values represent better performances. The dataset was bootstrapped 10^3^ times. Wide bars represent average values and error bars the 95% CI. B) Correlation between expression of the 109 Polycomb targets used in the model and the differentiation (binary) outcome for each of the two fates. The high correlation between fates (r=0.71) explains the fact that a model trained with PGCLC data can predict DN differentiation effectively. Experimental data from this study and Ref. [7].

Overall, we have demonstrated that the limited number of PcG targets identified in this study can predict differentiation efficiencies into two cell types with a high degree of accuracy. In addition, our study provides a mechanistic understanding of this transcriptional heterogeneity, which appears to be inherited along the differentiation trajectory towards pre-ME.

## Discussion

We have used a combination of experimental and computational techniques to uncover the chromatin-based mechanisms that underpin transcriptome differences between hiPSC lines. We have also evaluated the differentiation efficiencies of ten hiPSC lines for two different cell fates (PGCLCs and DEs), finding highly variable efficiencies among lines and fates. Each cell line was capable of differentiating into at least one of the target cell types (Fig. 1C and Supplementary Fig. 3C). By profiling the transcriptome and three different histone PTMs and integrating them with chromatin accessibility measurements, we obtained a unique epigenomic dataset. We analysed this dataset using an SVM-based computational pipeline to dissect the different regulatory modes. We conclude that *1)* the modes involving H3K27me3 displayed the most consistent differences (Fig. 6A, B), *2*) the genes in these modes were predictive of differentiation efficiencies (Fig. 7) and *3)* they had the capacity to create two distinct states, suggesting a digital form of histone PTM regulation (Fig. 4C). Crucially, this bimodality was tightly linked to the repression of the associated genes: not only did it divide the datapoints between high and low H3K27me3 levels, but also between higher and lower expression levels (Fig. 4D).

Furthermore, our mathematical modelling revealed that this digital mode of regulation can be explained by the read-write feedback of Polycomb complexes, a positive feedback loop that propagates the H3K27me3 mark and creates two discrete states (ON/OFF). To our knowledge, this is the first instance of genome-wide quantitative modelling of digital Polycomb regulation. Previous modelling studies concerned with digital regulation have either focused narrowly on a single locus [48–50], or performed genome-wide quantitative modelling of datasets without discriminating between unimodal and bimodal behaviour [21]. Nevertheless, digital regulation of chromatin has been shown for certain genes in yeast [51, 52], plants [53] and mouse neural progenitor cells [54]. Here, we demonstrate that a similar behaviour occurs in some genes in hiPSCs, and that it can impact transcriptional output.

Importantly, our mathematical model showed that even slight alterations to the parameters for the overall H3K27 methylation rates can cause dramatic changes in histone PTM levels. The fact that a small dysregulation of Polycomb silencing could cause such a marked effect in a subset of genes is consistent with the observation that the hiPSC lines in this study display highly similar genome-wide transcription and chromatin regulatory landscapes, with pronounced differences in both restricted to specific genomic loci. One potential candidate for this dysregulation is *EPOP*, which is expressed at higher levels in the cell lines with altered gene expression and chromatin profiles in H3K27me3 regulatory modes (Fig. 2D and Fig. 6A, C). EPOP is the only Polycomb accessory protein that was differentially expressed among the studied cell lines. EPOP has been shown to modestly reduce the binding of Polycomb to chromatin [55, 56], but, according to the model, this modest effect may be sufficient to completely switch the transcriptional regulation of specific target genes.

Finally, we showed that this dysregulation of Polycomb silencing is predictive of differentiation efficiencies for two cell types (PGCLCs and DNs). Our predictions greatly improve on past models in terms of performance metrics and in terms of biological interpretation, even though we used expression from only a small subset of PcG targets, rather than a more than two orders of magnitude larger genome wide set [7]. While the fact that expression from this subset was predictive does not imply direct causality, our analysis highlights the importance of locus-specific H3K27me3 in differentiation efficiency for both the neuronal and germ cell lineages. Further strengthening this link, we also demonstrated that transcriptional heterogeneity of H3K27me3 targets was propagated to an intermediate state in the differentiation trajectory (pre-ME, Fig 6D).

Altogether, this work has shown that hiPSC lines exhibit heterogeneous, fate-dependent differentiation efficiencies, which was linked to variability in H3K27me3 regulation. Taking into account the impact of repressive histone modifications on transcription, and the number of differentially expressed transcription factors among the hiPSC lines, we propose that the dysregulation of H3K27me3 levels at specific gene loci is one important factor that influences the diverging differentiation properties between hiPSC lines. However, we note that genetic variations between donors [9, 10], DNA methylation [5], or sex [57] have also been shown to contribute to these differences. Nevertheless, emerging evidence emphasising the role of the H3K27me3 histone mark in regulating pluripotency supports our findings [12, 58, 59], further reinforcing the hypothesis that histone modifications modulate the capacity of pluripotent stem cells to differentiate into target cell types.

## Methods

### hiPSC culture

The human induced pluripotent stem cell (hiPSC) lines used in this study (HPSI1113i-bima_1, HPSI0114i-kolf_2, HPSI0114i-kolf_3, HPSI0214i-kucg_2, HPSI0514i-letw_5, HPSI1113i-podx_1, HPSI1113i-qolg_1, HPSI0314i-sojd_3, HPSI0214i-wibj_2, and HPSI0215i-yoch_6) were purchased from the Human Induced Pluripotent Stem Cells Initiative (HipSci; https://www.hipsci.org/). All cell lines were deposited by the Wellcome Trust Sanger Institute (https://www.sanger.ac.uk/), and banked at European Collection of Authenticated Cell Cultures (ECACC). ECACC’s quality control testing methods are accredited in accordance with the recognized International Standard to ISO/IEC 17025:2005. All hiPSC lines were purchased from Culture Collections of the UK Health Security Agency (https://www.culturecollections.org.uk/) and tested negative for mycoplasma contamination.

hiPSCs were cultured on vitronectin-coated (Gibco, A14700) dishes in TeSR-E8 medium (Stem Cell Technologies, #05991). The medium was changed daily, and cells were passaged as clumps every 4-5 days using Versene (Gibco, 1504033) or 0.5mM EDTA solution (Life Technologies, AM9260G) in PBS. 10μM Y-27632 2HCl (Selleckchem, S1049) was added for 1 day after passage. All assays were performed between passages 4 and 8.

### Pre-ME differentiation

Induction of human PGCLCs was performed as previously described [22]. Briefly, confluent hiPSCs were washed once with 1x PBS (Gibco, 20012-019) and then trypsinised with 0.25% Trypsin-EDTA (Gibco, 25200056). The cells were then incubated at 37°C for 2.5-3 min and resuspended in MEF medium (DMEM/F-12 (Gibco, 21331-020), 10%FBS (Gibco, 10270106), 2mM L-Glutamine (Gibco, 25030081), 100 U/ml Penicillin-Streptomycin (Gibco, 15140122)) to stop the reaction. The cells were passed through a 35 μm cell strainer (Falcon, 352235) to remove the clumps and the resulting suspension was centrifuged at 200g for 4 min. The dissociated cells were then seeded in vitronectin-coated 12-well plate at a density of 200 000 cells/well. Cells were cultured in mesendoderm medium (aRB27 medium (Advanced RPMI 1640 (Gibco, 12633012), 1% B-27 supplement (Gibco, 17504001), 1x MEM NEAA (Gibco, 11140-035), 2mM L-Glutamine, 100 U/ml Penicillin-Streptomycin) supplemented with 100ng/mL Activin A, 3μM CHIR and 10 μM Y-27632 2HCl) for 12h.

### PGCLC differentiation

For PGCCL induction, dissociated Pre-ME cells were washed with PBS, trypsinised for 2,5-3 min at 37°C, resuspended in MEF, passed through a strainer, centrifuged at 200g for 4 min, and resuspended in PGCLC medium (aRB27, 500 ng/ml BMP4 (R&D Systems, 314-BP), 10 ng/ml hLIF (SCI), 100 ng/ml SCF (R&D Systems, 455-MC-010), 50 ng/ml mEGF (R&D Systems, 2028-EG-200), 10 μM Y-27632 2HCl). The cells were then seeded in ultra-low-attachment 96-well plates (Costar, 7007 or Greiner, 650979) at a density of 5000 cells/well and the resulting EBs were cultured for 4 days.

### DE differentiation

DE induction was performed as previously described [22]. Briefly, Pre-ME cells were cultured in ME medium for another 12 h (24 h in total), then the cells were washed once with PBS and the medium was changed to DE medium (aRB27, 100ng/mL Activin A, 0.5 μM LDN212-854). The DE cells were collected 2 days after the medium change.

### EB processing, cryosectioning and immunofluorescence

For PGCLC staining, EBs were collected and washed with PBS. For fixation, EBs were incubated in 4% (w/v) paraformaldehyde (Agar Scientific, AGR1026) for 20 min at room temperature. EBs were then washed three times in PBS and incubated in 10% sucrose (Carl Roth, 57-50-1) at 4°C for either 1 h or overnight, followed by incubation in 20% sucrose at 4°C for 1h. EBs were then embedded in OCT (Cell Path, 6478.2), incubated for 30 min at 4°C and frozen at -80°C until further use. For cryosectioning, a Leica cryostat was used, and 8μm sections of EBs were collected onto charged slides (Epredia, J7800AMNT) which were stored at -80°C until further use. Slides were first washed three times for 5 min in PBS and then incubated in permeabilisation buffer (PBS, 1% BSA (Carl Roth, 8076.2), 0.1% Triton X-100 (Thermo Fisher, BP151-100)) for 30 min in a humidified chamber. The slides were then incubated with primary antibodies (Supp. Table 2) overnight at 4°C. The next day, the slides were washed three times in PBS and then incubated with secondary antibodies (Supp. Table 2) for 1 h at room temperature. Slides were then washed twice in PB, incubated with DAPI and washed again with PB before mounting with Vectashield (H-1000, Vector Laboratories). Microscopy was performed using an LSM confocal microscope (Zeiss, LSM 980). For hiPSC or DE characterisation, the cells were grown on vitronectin-coated 8-well chamber slides (Ibidi, 80821) (hiPSCs) or on glass coverslips (DE cells) and the slides were processed as described above. The Object Scan plugin (https://github.com/orgs/gurdon-institute/repositories; [60]) was used in Fiji to quantify the images.

### Fluorescence-activated Cell Sorting (FACS)

FACS was performed as previously described with some minor modifications [22]. Briefly, human EBs were harvested on day 4, washed with PBS and incubated in 0.25% Trypsin/EDTA in a thermocycler at 850 rpm at 37°C for 5-12 min. The reaction was then stopped with 3% FBS in PBS and the cell suspension was pipetted until dissociated into single cells and then passed through the strainer. The cells were then centrifuged, and the cell pellet was resuspended in 3% FBS in PBS. Dissociated cells were then stained for EpCAM-APC (Biolegend, 324208) and ITGalpha6-BV421 (Biolegend, 313624) for 1 h in the dark on ice. Afterwards, the cells were washed in 3% FBS in PBS and subjected to FACS using SONY Cell Sorter SH800Z. FACS data were analysed using FlowJo software.

### RNA-seq

For each cell line, ∼ 1 million human iPS or pre-ME cells were prepared per biological replicate. Cells were harvested and washed in Dulbecco’s Phosphate Buffered Saline (DPBS, Cytiva HyClone, SH30028.02). Total RNA was purified using the Monarch Total RNA Miniprep Kit (NEB, T2010S). An aliquot of the total RNA was taken for integrity analysis using the Agilent RNA 6000 Pico Kit (Agilent, 5067-1513) on a 2100 Bioanalyzer (Agilent). The rest of the total RNA was stored at -80°C until library preparation. RNA-seq libraries were generated using the NEBNext Ultra II Directional RNA Library Prep Kit for Illumina (NEB, E7760L), NEBNext Poly(A) mRNA Magnetic Isolation Module (NEB, E7490L) and NEBNext Multiplex Oligos for Illumina (Index Primers Set 1, 2, 3 and 4, NEB, E7335S, E7500S, E7710S, E7730S), using 500 ng total RNA as starting material for each preparation, and 8 final PCR cycles to obtain sequencing-ready libraries. Library concentration and quality were analysed using the Agilent High Sensitivity DNA Kit (Agilent, 5067-4626) on a 2100 Bioanalyzer (Agilent). The RNA-seq libraries were sequenced in HiSeq2500-RapidRun 50bp Paired End runs on the HiSeq sequencing platform (Illumina).

### CUT&Tag

CUT&Tag was performed using the EpiCypher protocol as described previously^20^ with minor modifications. hiPSCs were washed once with DPBS (Life Technologies; 14190144) and incubated with Accutase (StemCell Technologies; 07922) for 5 min at 37°C. After centrifugation for (3 min at 300 x g at RT), the cells were resuspended in 1X DPBS and counted. 2 × 10^5^ cells per CUT&Tag reaction were centrifuged for 3 min at 300 x g at RT and then resuspended in 100 μl/sample of cold NE Buffer (20 mM HEPES–KOH, pH 7.9, 10 mM KCl, 0.1% Triton X-100, 20% Glycerol, 0.5 mM Spermidine (Sigma-Aldrich, 05292), 1x Roche cOmplete^TM^, Mini, EDTA-free Protease Inhibitor) and incubated for 10 min on ice. After centrifugation (3 min at 600 x g at RT), the nuclei were resuspended in cold NE Buffer to achieve a final concentration of 1×10^6^ nuclei/ml. 1 × 10^5^ nuclei per reaction were subjected to CUT&Tag. A total volume of concanavalin A (ConA) beads (EpiCypher, 21-1401) of 11 μl per sample was transferred to a 1.5 ml tube for batch processing. The ConA beads were washed twice on a magnetic stand with 100 μl/sample Bead Activation Buffer (20 mM HEPES, pH 7.9, 10 mM KCl, 1 mM CaCl_2_, 1 mM MnCl_2_) and resuspended in a final volume of 11 μl per sample. 10 μl beads were aliquoted into separate tubes for each sample and incubated with 100 μl of nuclei for 10 min at RT. The tubes were placed on a magnet separator (DynaMag-PCR; ThermoFisher Scientific), and the supernatant was discarded. The beads were then resuspended in 50 μl of cold Antibody150 Buffer (20 mM HEPES, pH 7.5, 150 mM NaCl, 0.5 mM Spermidine, 1x Roche cOmplete^TM^, Mini, EDTA-free Protease Inhibitor, 0.01% digitonin (Sigma-Aldrich, D141), 2mM EDTA) containing primary antibody in a 1:50 dilution (anti-H3K9me3, Active Motif 39062; anti-H3K27me3, Active Motif; 39157; dilution 1:50; anti-H3K4me3, Active Motif; 39160). Samples were incubated overnight at 4°C on an orbital shaker. On the following day, the tubes were placed on a magnet separator, and the supernatant was discarded. The beads were incubated in Digitonin150 Buffer (20 mM HEPES, pH 7.5, 150 mM NaCl, 0.5 mM Spermidine, 1x Roche cOmplete^TM^, Mini, EDTA-free Protease Inhibitor, 0.01% digitonin) containing 0.5 μg of secondary antibody (anti-rabbit Epicypher; 13-0047) for 1 hour at RT. The beads were then washed twice with Digitonin150 Buffer and then resuspended in 50 μl of Digitonin300 Buffer (20 mM HEPES, pH 7.5, 300 mM NaCl, 0.5 mM Spermidine, 1x Roche cOmplete^TM^, Mini, EDTA-free Protease Inhibitor, 0.01% digitonin), and 1.25 μl of CUTANA pAG-TN5 (EpiCypher, 15-1017) was added. After incubation at RT for 1 hour, the tubes were placed on a magnet and washed twice with Digitonin300 Buffer.

The beads were then resuspended in 50 μl of chilled Tagmentation Buffer (Digitonin300 Buffer, 10 mM MgCl_2_) and incubated for 1 hour at 37°C in thermocycler. The tube was placed on a magnetic separator and the supernatant was discarded. The beads were washed once with 50 μl RT TAPS Buffer (10 mM TAPS pH 8.5, 0.2 mM EDTA). The beads were resuspended in 5 μl of RT SDS Release Buffer (10 mM TAPS, pH 8.5, 0.1% SDS) and vortexed at maximum speed for 10 seconds followed by a brief centrifugation. The beads were then incubated for 1 hour at 58°C in a thermocycler. Following incubation, 15 μl of RT SDS Quench Buffer (0.67% Triton-X 100 in molecular grade H_2_O) was added to the beads and vortexed at maximum speed for 20 seconds, followed by a brief centrifugation.

For library amplification, 1 μl of each barcoded i5 and i7 primers (10μM stock) was used and PCR was performed using CUTANA High Fidelity PCR mix (EpiCypher, 15-1018), using the following PCR programme: 58°C for 5 min, 72°C for 5 min, 98°C for 45 s, 11-13 cycles of 98°C for 15 s, 60°C for 10 s, 72°C for 1 min. Libraries were purified twice using 1X volume of AMPure XP Beads (Beckman Coulter; A63881) and resuspended in 15 μl of 0.1X TE buffer. Following quality control using the Agilent High Sensitivity DNA Kit (Agilent, 5067-4626) on a 2100 Bioanalyzer (Agilent), the CUT&Tag libraries were multiplexed and sequenced (paired-end 150 bp) on a NovaSeq 6000 instrument (Illumina).

### Sequencing data processing

Raw RNA-seq reads were assessed for quality using FastQC v0.11.8 (www.bioinformatics.babraham.ac.uk/projects/fastqc). Reads were aligned to the human reference genome GRCh38 with STAR v2.5.2b. PCR duplicates were removed using Picard v2.9.0 (https://broadinstitute.github.io/picard/). Gene-level quantification was performed using HTSeq-count v0.10.

CUT&Tag data were processed and aligned to the GRCh38 reference genome as previously described [29]. After alignment, reads with mapping quality below 30 were filtered out using SAMtools v1.9. PCR duplicates were removed using Picard.

### Data

The data used in this work included the transcriptome of 10 iPSC cell lines, obtained by RNA-seq. We obtained transcript counts for all ten cell lines (two independent replicates for each cell line and three in the case of Yoch6). We focused our efforts on protein-coding genes of autosomes (to avoid confounding effects that may take place in sex chromosomes). The input for the differential gene expression analysis are the raw counts per gene. For every other analysis of the transcriptome, the magnitude used was transcripts per million (TPMs), since this metric normalises by the length of each gene, making it dimensionless and more reliable [61].

In terms of the epigenome, CUT&Tag was performed for the histone marks H3K27me3, H3K4me3 and H3K9me3 in all these lines, with two replicates for each. In addition, results from ATAC-seq assays were obtained from Ref. [30]. We used the bigwig files (normalised to reads per genomic content, i.e. 1×genomic coverage) from independent replicates to establish the enrichment of a given epigenetic mark above the genome-wide average (area under the curve) at specific genomic features, namely: genes (from gene start to gene end), promoters (defined as between 1 kbp upstream and downstream of the gene start) and gene bodies (the entirety of the gene except for the first kbp). Finally, we also used transcript counts at the transcript level to identify the most used transcription start site (TSS) in each gene. Then, these TSSs were also used to quantify enrichment in a window of 1 kbp upstream and downstream from it, similar to the procedure for promoters. These are the four types of enrichment quantifications used in the analysis below. In the case of the dataset used for the SVM, we quantified the enrichment using each of these four annotations but considering the merged bigwig file from all independent replicates. Where appropriate, we used the ENSEMBL annotation for the human genome (hg38, release 110).

### PCA

For the PCA analysis of the transcriptome, TPM values were normalised to unit variance (so that every gene has similar importance) and then used to plot the PCA diagram, with the coordinates of each datapoint being specified by the normalised gene TPMs for a given replicate in a particular cell line. Similar procedures were used to plot the PCA diagrams of histone marks and accessibility assays. The only difference, in these cases, is that we used the enrichment quantifications for each genomic feature (separately) obtained from the epigenomic assays instead of TPM values.

### Transcriptome scatter plots

For the transcriptome scatterplots we took the TPM values of each cell line and computed the logarithm (base 2) of the TPM value and added a constant to avoid taking the logarithm of 0, for cases with zero counts. We plotted on the X axis the results for one cell line and in the Y axis the results for another one, and we computed the Pearson correlation in between the two datasets. We note that the resulting Pearson correlation between cell lines is quite sensitive to the value of the constant added: the smaller this value, the smaller the correlation. Unlike in other studies, here we chose a smaller value for this constant (0.01).

### Differential expression analysis

We have data for 10 cell lines, and we are interested in any difference between any pair of cell lines. Thus, we need to perform multiple pairwise comparisons between all cell lines, to obtain a final list of differentially expressed genes (DEGs) that will be flagged for further analysis. When performing multiple pairwise comparisons the resulting threshold p-value for statistical significance needs to be adjusted (decreased) to take into account the larger false discovery rate of performing multiple pairwise comparisons. For this, we used a Benjamini-Hochberg (BH) adjustment method to increase the p-value (to account for the larger false discovery rate).

Usually log2-fold changes (L2FCs) are also applied, to ensure the genes tagged as differentially expressed show substantial changes in transcription. However, the same adjustment procedure to control for multiple pairwise comparisons is not available for L2FCs. Instead, with the aim of reducing the false discovery rate, we chose a high threshold for L2FCs. The thresholds we used were 0.01 for the BH-adjusted p-values and 3 for the L2FCs. Therefore, for a gene to be considered as differentially expressed, there must be at least one pairwise comparison where both the adjusted p-value is below 0.01 and the L2FC is above 3. Additionally, we filtered out genes that had less than three samples (out of 21) with more than 10 counts. With these thresholds and filters, we identified 712 strongly DEGs. For this analysis, we used R (v 4.4.2) and DESeq2 [62] (v 1.46.0).

### Gene ontology analysis

We performed a gene ontology analysis for the DEGs. We used the clusterProfiler package (v 3.0.4) and enrichGO function [63] on the 700 DEGs. The universe, or background genes against which to test the enrichment hypothesis, is the same set of genes that pass the filtering for differential expression analysis, that is, around 15,000. The p-value adjusting method was also the BH method and we tested both the molecular function and the biological process ontologies.

### Comparison between hiPSC lines and hESCs

As a measure of similarity between hiPSC lines and hESC lines, we computed the Euclidean distance between the ten iPSC lines analysed here and an hESC line (H9, from the ENCODE database [47], experiment ENCSR712BRU). We took the Euclidean distances between transcriptomes in terms of the z-score. Z-score standardises means and variances for each gene independently (considering the variability among hiPSC and hESC lines), which implies that the measure is not sensitive to basal rates (as the correlation is) and it focuses only on differences. We normalised the distances by a factor of √*N*, where *N* is the number of genes considered in each instance, to control for the increase of dimensionality and maintain a stabilised metric across analyses.

### Likelihood of random overlap between lines with altered H3K27me3 regulation and differentiation efficiency

Given that four cell lines had low PGCLC differentiation efficiency and three cell lines displayed altered transcription and H3K27me3 regulation, we wanted to know the probability that, if three cell lines are chosen at random, these three will be part of the four cell lines with impaired PGCLC differentiation. If all groups of three are equally random, we just need to count the number of cases where groups of three would fall within a group of four (4 choose 3) and divide by the total number of groups of three we can create with ten cell lines (10 choose 3). Then, the probability is:

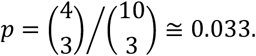

### Support Vector Machine analysis

SVMs are supervised classification methods, which, in their simplest form and given a set of training datapoints (each annotated with one of two possible classes), produce a hyperplane that separates the datapoints according to their class, maximising the distance from the closest datapoints to the hyperplane [64]. This technique enables two outcomes: to predict whether a gene will be highly or lowly expressed from its histone marks/accessibility, and to classify the different genes according to their regulatory mechanism. In addition, our dataset (10 very similar cell lines with transcriptional and epigenomic information) is well suited for this approach since it enables us to analyse each DEG separately (with 10 datapoints per gene), allowing us to capture the specific characteristics of each target. 10 datapoints per gene, in a four-dimensional space (accessibility and three histone marks), would usually be too small a dataset, but the robustness of SVMs nevertheless enables this approach with satisfactory precision. The procedure outlined below was performed on a per-gene basis.

### Binarization of expression

The starting point is a TPM value per DEG per cell line (averaged across replicates). For each gene, its TPM values across cell lines were binarized, that is, whether this specific gene is highly or lowly expressed in each cell line. To that end, we used a K-means algorithm [65] to cluster those 10 TPM values into zeroes and ones. To get the best description of the data, we employed both logarithmic and linear scales, i.e., we performed the K-means clustering for the TPM values in both logarithmic scale and linear scale. To decide which of the scales should be used in each case, we chose the better performing one according to its inertia (the sum of squares of the data in each cluster to the centre of the cluster). Thus, we can compare the goodness of the clustering in between the two scales by computing, for each scale, the ratio between the inertia when clustering into two clusters over the inertia when clustering into a single cluster, as a measure of how much better the two clusters (high and low expression) are with respect to a single cluster. If this ratio is smaller in one scale than in the other, it implies that the clustering is more accurate in that first scale. In general, this clustering is performed using a logarithmic scale since changes are more pronounced in terms of fold-changes than absolute differences. However, some genes have TPMs of zero for some cell lines, which makes the logarithmic scale invalid and, thus, we use the linear scale for these genes. Finally, it is worth noting that, despite this binary classification being a simplification of reality, it produces a reasonable classification that can be fed into the SVM, resulting in high accuracy predictions (see below).

### SVM description

We built a support vector machine (SVM) that classifies RNA levels (high versus low expression) based on histone marks and accessibility data. For each gene and each cell line, we fed the SVM a value of H3K4me3, H3K9me3, H3K27me3 and ATAC (see main text). For accessibility and H3K4me3 data, given that these profiles display much more pronounced and narrow peaks, we took the annotation that produced the maximum value for each gene among the following three: 1 kbp upstream and downstream of the gene start, 1 kbp upstream and downstream of the most used TSS, and the average over the whole gene. Usually, the whole gene average is the smallest metric, but we included it for cases where we failed to find the peak at either the gene start or TSS. This input data, together with the binarized RNA expression metric, is enough to train a SVM for each gene independently. For added robustness, it is convenient to normalise all of the input data to values between zero and one. Thus, if *x*^*j*^ is the enrichment of the *k*-th histone mark (or ATAC) around gene *i* for cell line *j*, the normalised quantity that is fed into the SVM is given by

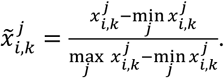

We define the scale factor as

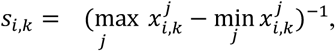

for later convenience. Once the SVM (linear kernel) is trained with the normalised data, the result is a hyperplane that separates highly expressed from lowly expressed cell lines in a four-dimensional space for the normalised values of the epigenomic variables at the gene of interest for each cell line. This is the first outcome of the SVM: a classification method to predict high/low gene expression based on its epigenetic marks/accessibility. In terms of parameters, we used a linear kernel and a high *C* parameter (*C* = 100). The *C* parameter controls the strength of the penalty for misclassification. A low *C* parameter would try to maximise the margin (distance between the hyperplane and the nearest point) instead of maximising classification accuracy. For an example of the application of the SVM, see the Supplementary Materials.

### SVM as a prediction tool: binary classification

To test the performance of the SVM (and the entire analysis pipeline, from DEG identification to SVM training, passing through transcription binarization) as a predictor of transcription, we included 9 cell lines in the training set and sought to predict the transcription of the 10^th^ (e.g., Qolg1) based on its epigenomic variables and the trained SVM. In this case, the binarization of expression is performed with the nine cell lines in the training data set and whether the 10^th^ cell line is high or low expression (ground truth for this test) will depend on which cluster centre its expression is closest to. In addition, a calibration was performed by binning into 10 groups the datapoints and predictions to compare the predicted probability of the SVM (using Platt scaling, see below) to the actual fraction of positives in the group (Supplementary Fig. 5D).

### Comparison with logistic regression

For benchmarking purposes, we sought to compare our SVMs with a more standard logistic regression approach, since it is an efficient linear classification method. The comparison between algorithms was performed in terms of the overall accuracy and the area under the receiver operating characteristic (AUROC) curve. The ROC curve links the true positive and false positive rates for each threshold in the binary classification and the higher the AUROC (bounded from above by 1), the more accurate the classifier.

To obtain the ROC curve, the SVM is required to produce scores (rather than binary predictions), which can be interpreted as the probability of a prediction. To that end, we used Platt scaling [34] to fit the output of the SVM to a sigmoidal curve that represents a probability function, which is then evaluated at the test datapoints to obtain a probability score for each prediction. Usually, the same SVM (same parameters for the hyperplane and fit to sigmoidal) would be used to predict multiple datapoints, resulting in a vector of scores (one for each datapoint), from which we can obtain the ROC curve. In this case, due to the limited datapoints per gene, we decided to train a SVM in each gene with 9 datapoints and seek to predict the 10^th^. Therefore, to obtain a vector of scores for the ROC curve, we independently trained the SVMs for each gene (as described above) and predicted a score for the 10^th^ datapoint in each gene. We obtained a vector of scores by pooling all genes together.

With the vector of scores from our classifier (and a vector of ground truths for the test dataset) we can obtain the ROC curve by varying the score threshold (from 0 to 1) above which a prediction would be considered positive. Collectively, this set of points (one for each different threshold) in the true positive rate vs false positive rate space makes the ROC curve. It should be noted that, given that we are pooling scores from different genes and SVMs together, the ROC curve we are plotting is not the usual ROC curve, obtained from evaluating many datapoints with the same SVM. However, it still serves as a collective measure of the goodness of classification of the overall procedure.

To obtain the ROC curve for the logistic regression we followed an analogous procedure as with the SVM. When evaluating the accuracy and AUROC of SVMs or logistic regressions, we bootstrapped the training dataset 10^3^ times, to obtain confidence intervals for performance when predicting each cell line’s expression patterns.

### SVM as an explanatory tool: coefficients and clustering

For the second outcome (identification of regulatory mechanisms), we leverage the coefficients of the hyperplane defined by the SVM procedure. The coefficients of the hyperplane define the vector normal to the surface, which in this case and given our conventions, is the direction of upregulation of transcription. Thus, in our four-dimensional case, for each gene *i*, we obtain ***n***_*i*_ = (*n*_*i*,*K*4_, *n*_*i*,*K*9_, *n*_*i*,*K*27_, *n*_*i*,*ATAC*_). The combination of the hyperplane coefficients (how epigenetic marks/accessibility relate to transcription) and the dynamic ranges (how much variability each epigenetic mark/accessibility has in each gene) can be very informative for the regulatory logic, but, in their raw form, the coefficients of the SVM analysis cannot be directly used and need to undergo some preprocessing.

We need a quantity that incorporates both the directionality information of the SVM hyperplane, combined with the dynamical range of the original data. The inclusion of the dynamical range is important to gauge the strength of the effect, e.g., a dynamical range just above the genome-wide mean for that mark is unlikely to have a strong effect in transcription, even if, due to some spurious correlation/noise, the SVM normal vector points in that direction. To that end, first, each of the four coefficients that define the hyperplane was normalised using the summed square of the four coefficients. This allows us to ascertain the relative contribution of each variable and permits an interpretation as a vector within a sphere of radius one. The normalised hyperplane coefficients are:

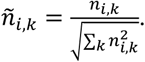

Second, the information lost when normalising to feed epigenomic data to the SVM needs to be recovered. To that end, we take the logarithm of the inverse of the scale factor *s*_*i*,*k*_ (if *s*_*i*,*k*_ > 1 we take the log to be zero to avoid negative results). If *s*_*i*,*k*_ ≥ 1, the dynamical range is the same as the genome-wide mean or smaller (due to the normalisation of the raw data), implying that any changes in that mark, at that gene, are relatively small and we therefore neglect such changes. This leads to the quantity we are looking for, the log-rescaled coefficient:

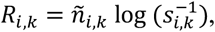

which combines the information of the SVM with the dynamical ranges in logarithmic scale, while preserving the signs of the SVM vector. Note that, while the epigenomic data was normalised by *s*_*i*,*k*_ before feeding it to the SVM, the hyperplanes coefficients had not (i.e., this is not directly undoing an earlier operation). Finally, to avoid biasing our analysis towards quantities that (due to their own nature) show greater dynamical ranges, we normalise the maximum of log-rescaled coefficients across all genes, for each histone mark/accessibility assay:

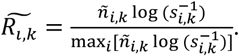

The rationale for using the logarithm of the normalisation constant is that we believe that fold changes are more instructive than absolute differences. The reason for using the normalised hyperplane coefficients rather than their logarithm is that the hyperplane coefficients might be negative, so cannot themselves be recast on a logarithmic scale. In addition, the signs of the hyperplane coefficients should be kept, since they specify whether a given epigenetic mark is activating (positive) or repressing (negative) with respect to transcription.

After the processing of the SVM outputs, we can now proceed to clustering DEGs into clusters according to their regulatory logic. To that end, we use again a K-means algorithm for clustering the log-rescaled coefficients 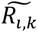. We clustered the genes into 2-20 clusters (1000 independent random initialisations) and examined different measures for the goodness of clustering (Silhouette score, Calinski-Harabasz score and Davies-Bouldin score). The Calinski-Harabasz score identified 4 as the optimal number of clusters, while the Davies-Bouldin and Silhouette scores identified 5 as the optimal cluster number (see Supplementary Fig. 5A). However, the Silhouette and Davies-Bouldin scores also had another local optimum for 10 or 11 clusters, with similar scores as for the cluster of 5. Given that clustering into only 4 or 5 clusters is too coarse for the downstream analyses we are aiming for, we chose 10 as the number of clusters to keep.

### Z-score RNA cluster plot

We plotted the RNA TPM values of the DEGs in each cluster for all 10 cell lines. Using Seaborn’s clustermap function, the genes and cell lines were hierarchically clustered and plotted (with the mean expression of each gene set to 0 and the variance normalised to 1, i.e., using the z-score).

### Bimodality

To ascertain whether the datasets we are working with are bimodal in the different regulatory contexts, we computed the Hartingan’s dip statistic (HDS) for each histone mark in each of the different clusters. The HDS measures the distance from the empirical distribution to the closest unimodal distribution. Thus, the higher the statistic, the further away it is from a unimodal distribution. To compute an empirical HDS for each cluster (set of genes with varying characteristics, such as length or CG content), it is necessary to normalise the dynamical range of each variable at each gene to the [0,1] interval. When normalising each gene into the [0,1] interval, there will be one value that is mapped to 0 and one that is mapped to 1 (out of a total of 10 values). When pooling together dozens of genes, this results in an artificial over-representation of 0s and 1s that could bias the results of bimodality metrics. To avoid this issue, for each gene, one 0 value and one 1 value is dropped from the dataset (in total, 2 values per gene are dropped). Pooling the resulting values within each cluster, we obtain an empirical distribution for which an HDS value can be obtained. We note that this procedure (since it discards the extremes for each gene) results in a conservative metric for bimodality.

The scripts in this section were written and executed in Python 3.10.11, using scikit-learn [65] version 1.4.1, pandas version is 2.2.1, numpy version is 1.26.2, scipy version 1.12.0 and seaborn version is 0.13.2.

### Mathematical model

The model describes the fraction of lysine residues 4 and 27 of histone 3 that are methylated in two different regions: promoter (±1kbp around the gene start) and gene body (from +1kbp of the gene start to the gene end). The definitions of these annotations follow those of the main text and the computational analysis of experimental data.

For lysine residue 27, since it has been shown to be important for theoretical modelling, we consider four states: H3K27me0, H3K27me1, H3K27me2 and H3K27me3. Methylation of the H3K27 residue is possible both at the promoter and gene body (variables for promoter and gene body are distinguished by a subscript, *p* or *b*, respectively). For lysine residue 4, we describe two states: either unmodified (H3K4me0) or trimethylated (H3K4me3). We only consider two states for H3K4, since there is little evidence for any feedback in these states and methylation at this residue occurs on a faster timescale [66]. We assume methylation of residue H3K4 is only possible at promoter regions, as H3K4me3 in the gene body is usually negligible. In addition, we assume that methylation at H3K4 and at H3K27 are mutually exclusive, since they cannot take place on the same histone tail [67].

The fractions of histones with each modification are represented by [·] and their quantitative dynamics are given by:

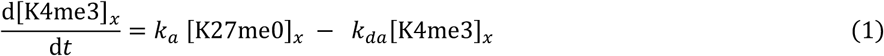

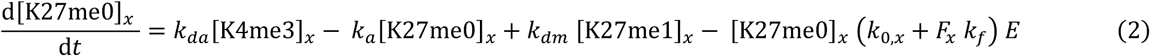

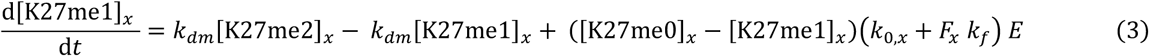

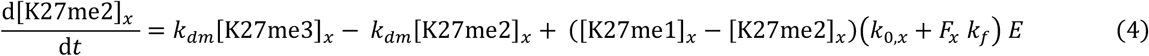

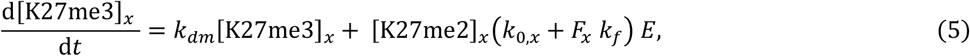

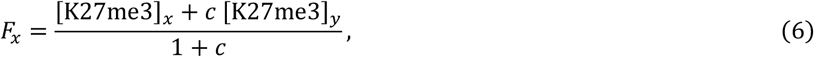

*x,y* ∈ (*p*,*b*) and *x* ≠*y*. Thus, the only coupling between the two regions is due to the feedback term *F*, parametrised by *c* (which is always smaller than 1, ensuring that these regions interact more within themselves than with each other). The construction of the model reflects the fact that if interactions are stronger with the other region they have to be weaker within the same region, as a locus can make a limited number of contacts. In addition, the cell cycle is also simulated, by halving all histone marks by a factor of 2 (and increasing consequently the fraction of H3K27me0 histones) every period *T*.

The H3K27 demethylation rates are parametrised by *k*_*dm*_. *k*_0,*x*_ represents the strength of the basal rate at region *x*, and *k*_*f*_, that of the feedback rate. *k*_*a*_is the rate of H3K4 methylation, and *k*_*a*_ ≠ 0 only at the promoter region. *k*_*da*_is the rate of H3K4me3 demethylation. The parameter *E* is cell-line dependent. Therefore, in order to describe the different configuration in which a gene from any of the Polycomb clusters can be found, the parameter *E* will be varied.

Finally, similar to previous models [20], we assumed that the transcriptional response takes the form:

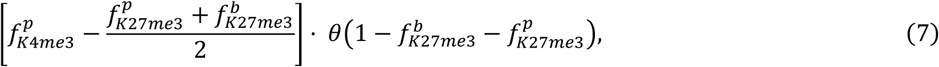

provided that 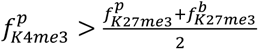 (otherwise the transcriptional output is 0). In Eq. (7), *θ* is the Heaviside (step) function and *f* denotes the normalised and noise-added outputs of the simulations (for details see below).

### Generation of synthetic data

First, we obtained parameters for a deterministic simulation, based on the underlying parameters for the cluster (see Table 1). For each cluster of genes, we obtained as many sets of random parameters as genes in the cluster. The random parameters were normally distributed, with mean at the cluster values and with a standard deviation of 10% of the mean value.

For each of these parameter sets, we obtained 10 datapoints (since we have 10 cell lines), by changing the parameter *E*. Thus, the simulations are deterministic once the parameters are fixed. The simulations are run (according to the equations of Methods—*Mathematical model*, starting from a silent—high H3K27me3—configuration) for 20 cell cycles to equilibrate, and sampled 100 times during the 21st cell cycle. The average (for each variable) of the samples in the last cell cycle is the only value stored from each simulation. This choice reflects the fact that, in experiments, cell cycles within the population were not synchronised and, hence, their outcome is an average of cells at different points of the cell cycle.

To simulate the spread of the data more realistically (simulating experimental noise), noise was added to the output obtained from each simulation independently. The random variables for this noise were drawn from a normal distribution (with mean 0.1 and variance 0.05) and added to the histone mark fractions (naturally bounded from above by 1). We took absolute values of this sum, for the unlikely event that it results on a negative value.

The 10 noisy datapoints that are the output of the simulation are then normalised between 0 and 1 (exactly the same procedure as described in the methods for the experimental data, to make them comparable). This procedure yields normalised outputs *f*, which are used to compute the transcriptional output, as per Eq. (7). Finally, we also add noise to this transcriptional output, by adding a normally distributed variable (again, with mean 0.1 and variance 0.05) and taking absolute values. Transcriptional output is also resized to span the [0,1] segment (as was done for the experimental data).

From here, any further downstream quantification (e.g., HDS) is made to compare with experimental data and, thus, follows exactly the same procedure as described in the methodology for the analysis of the experimental data.

### Parameter

We set *k*_*da*_ = 2 hr^−1^, to reflect the more rapid turnover of active marks (compared with the K27 demethylation rate *k*_*dm*_ = 1 hr^−1^). To simulate the cell-line specific differences, the parameter *E* is varied from 1 to 0.8 (0.8 representing weaker Polycomb activity, e.g., *Yoch6* cell line). As stated in the main text, we assume that *k*_0,*p*_ = 1.5*k*_0,*b*_ in all three clusters. All of the fixed parameters of the model are given in Table 1.

### Prediction of differentiation properties

We built a logistic regression model to predict the ability of hiPSC lines to differentiate into PGCLCs, based on the expression of 152 PcG targets. Ability to differentiate into PGCLCs was considered a binary variable: cell lines with low or null efficiency for PGCLCs, as per Fig. 1C, were considered as failing to differentiate, and the remaining cell lines as successful. The transcriptome in the training datasets (in TPMs) was normalised to the [0,1] segment across cell lines, for each gene independently. Five of these genes were excluded for consistency, since they did not appear in the expression processed files from the HipSci consortium (which used GRChg37 for the alignment).

To prevent overfitting an aggressive L2 regularisation was applied to the model (C=0.1, scikit-learn [65] version 1.4.1 LogisticRegression function). To further simplify the model, the training of the model was bootstrapped 10^4^ times (with C=1, to allow for a greater range of coefficients), and only features whose coefficient’s 95% CI did not overlap with 0 were retained, ensuring that the features were robustly predictive of differentiation. This resulted in 109 features remaining. When validating the PGCLC model, it was trained with these 109 features in 81% of the data (17 out of 21 RNA samples) and tested in the remaining 19%, with the entire dataset bootstrapped 10^3^ times.

To test the model in a context of DN differentiation, we downloaded RNA expression data (TPMs) for 143 hiPSC lines from the HipSci consortium (for details on the lines and files used, see Supplementary File 3); lines that had been assessed for neuronal differentiation [7]. Expression normalisation was performed in the same way, but independently from the expression data used in the PGCLC model. For the version of the model trained in neuronal data, the same procedure as for PGCLCs was followed, resulting in only 47 highly predictive PcG targets. Validation was performed using leave-one-out cross-validation, as done in Ref. [7], both for the PGCLC model (when applied to neuronal data) and for the model trained in neuronal data, all of them bootstrapped 10^3^ times for CI estimation.

## Data and code availability

Data will be made publicly available upon publication and the code used to perform the SVM analysis and mathematical model is available at https://github.com/AMovillaMiangolarra/hiPSCchromatin-SVM

## Acknowledgements

AMM and MH thank the BBSRC Institute Strategic Programme (BB/P013511/1) and the Wellcome Human Developmental Biology Initiative for funding. LL thanks the Biochemical Society for funding. UG was supported by a Sofja Kovalevskaja Award of the Humboldt foundation. OD was supported by the International Max Planck Research School for Genome Science. JK was supported by the International Max Planck Research School for Molecular Biology. SS was supported by a UKRI MRC Rutherford Fund Fellowship (MR/T016787/1). FT, YC, and SS were supported by the BBSRC Babraham Institute’s Epigenetics Strategic Programme Grant (BBS/E/B/000C0421) and a BBSRC Flexible Talent and Mobility Account (FTMA) 3 grant.

## Author Contributions

AMM and MH led the computational analyses and mathematical modelling, with support from LL. JK carried out iPSC culture and differentiation into PGCLCs and definitive endoderm, and validated differentiation efficiencies by microscopy and FACS analyses. FT cultured iPSCs and generated CUT&Tag libraries. OD carried out initial data processing and mapping, and YC generated RNA-seq libraries. UG and SS planned and supervised the experiments. UG, SS and MH conceived the project and secured funding. AMM, UG, SS and MH interpreted the data and wrote the manuscript, with input from all authors.

