## Supplementary materials, tables and figures for "Signatures of digital Polycomb regulation in functional iPSC heterogeneity between individuals"

### Supplementary Materials for “Signatures of digital Polycomb regulation in functional iPSC heterogeneity between individuals”

#### Supplementary Table 1

| Cell line | Donor ID | HipSci iPSC line | Sex | Age of donor | Ethnicity |
|---|---|---|---|---|---|
| ***Bima1*** | HPSI-bima | HPSI1113i-bima_1 | Male | 40-44 | White British |
| ***Kolf2*** | HPSI-kolf | HPSI0114i-kolf_2 | Male | 55-59 | White British |
| ***Kolf3*** | HPSI-kolf | HPSI0114i-kolf_3 | Male | 55-59 | White British |
| ***Kucg2*** | HPSI-kucg | HPSI0214i-kucg_2 | Male | 65-69 | White British |
| ***Letw5*** | HPSI-letw | HPSI0514i-letw_5 | Male | 70-74 | White British |
| ***Podx1*** | HPSI-podx | HPSI1113i-podx_1 | Female | 65-69 | White British |
| ***Qolg1*** | HPSI-qolg | HPSI1113i-qolg_1 | Male | 35-39 | White British |
| ***Sojd3*** | HPSI-sojd | HPSI0314i-sojd_3 | Female | 45-49 | White - Other |
| ***Wibj2*** | HPSI-wibj | HPSI0214i-wibj_2 | Female | 55-59 | White British |
| ***Yoch6*** | HPSI-yoch | HPSI0215i-yoch_6 | Female | 60-64 | White British |

#### Supplementary Table 2

| | Antibody | Company | Catalog # | Dilution |
|---|---|---|---|---|
| **Primary antibodies** | AP2γ | Santa Cruz | sc-12762 | 1:50 |
| | BLIMP1 | Cell Signalling | 9115 | 1:100 |
| | FOXA2 | Cell Signalling | 8186 | 1:500 |
| | SOX17 | R&D Systems | AF1924 | 1:750 |
| **Secondary antibodies** | Alexa647-donkey-α-mouse IgG | Thermo Fisher | A32787 | 1:400 |
| | Alexa488-donkey-α-goat IgG | Thermo Fisher | A11055 | 1:400 |
| | Alexa555-donkey-α-goat IgG | Thermo Fisher | A31572 | 1:400 |

### SVM example

We take the gene *LHX5* to illustrate the SVM procedure. For simplicity and easier visualisation, we will train the SVM with only H3K4me3 and H3K27me3 data (in the main workflow of this paper the full four-dimensional form of the SVM is used). For the *LHX5* gene, the raw data for the bivalent marks is shown in Fig. E1 A.

The first step is the binarization of transcriptional output. As stated in the methods section, we carry out the binarization procedure with a K-means algorithm for two clusters (high and low expression), both in linear scale and in logarithmic scale, and choose the optimal one. The metric we use as 'goodness of binarization' is the ratio between the inertia of the clustering into two classes over the inertia of a single class (computed for each scale independently). The lower the ratio, the more effective the binarization has been. In this case, we obtained a ratio of inertia in the linear scale of 0.35, while the ratio of inertia for the logarithmic scale was 0.07. Thus, we chose the logarithmic scale binarization. Additionally, and as explained above, we normalised the input data of both histone marks to the segment [0,1]. The resulting pre-processed data is shown in Fig. E1 B.

Now, we are in a position to train the SVM, which provides the line (we are only working in 2D in this example) that separates high expression from low expression cell lines and, at the same time, maximises the margin. The output of the SVM is shown in Fig. E1 C, where the output of the SVM is marked in black (both the line separating high/low expression and the normal vector ***n***). Therefore, if we wanted to predict transcription of an eleventh cell line, based on the enrichment of H3K4me3 and H3K27me3 at the *LHX5* locus, we would need to evaluate the enrichment levels of these marks to ascertain on which side of the line they fall. Moreover, the vector normal to the line, ***n***, informs us of how each of the histone marks are related to transcription. However, in order to systematically obtain insights regarding the regulatory logic at this locus, we need to post-process the output of the SVM and the original data, as explained above (see Methods—*SVM as an explanatory tool: coefficients and clustering*).

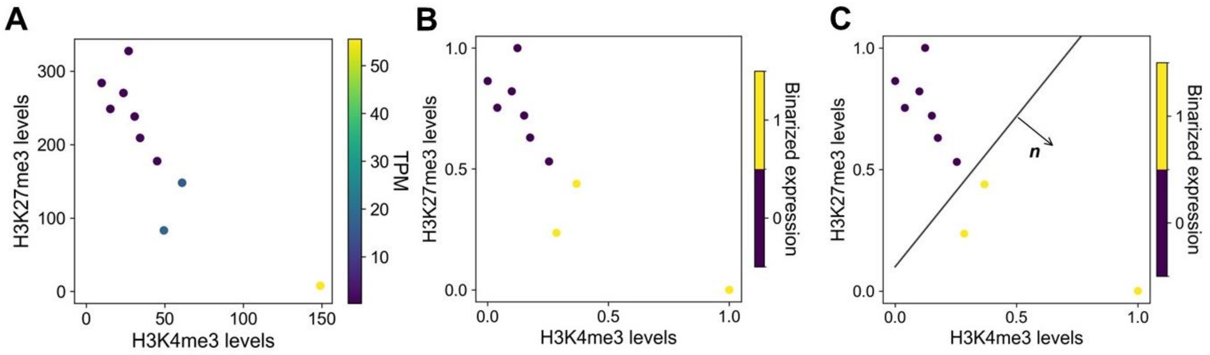

***Figure E1***. *Example of the SVM procedure. A) Raw data for the bivalent marks and transcription at the* LHX5 *gene. B) Same as A) but with normalised histone mark levels and RNA expression binarized. C) Output of the SVM, in terms of a line separating lower expressed from higher expressed cell lines.* ***n*** *is the vector normal to the line, which points in the direction histone marks need to change in order to upregulate transcription and is indicative of the type of epigenetic regulation in that gene.*

### Supplementary Figure 1

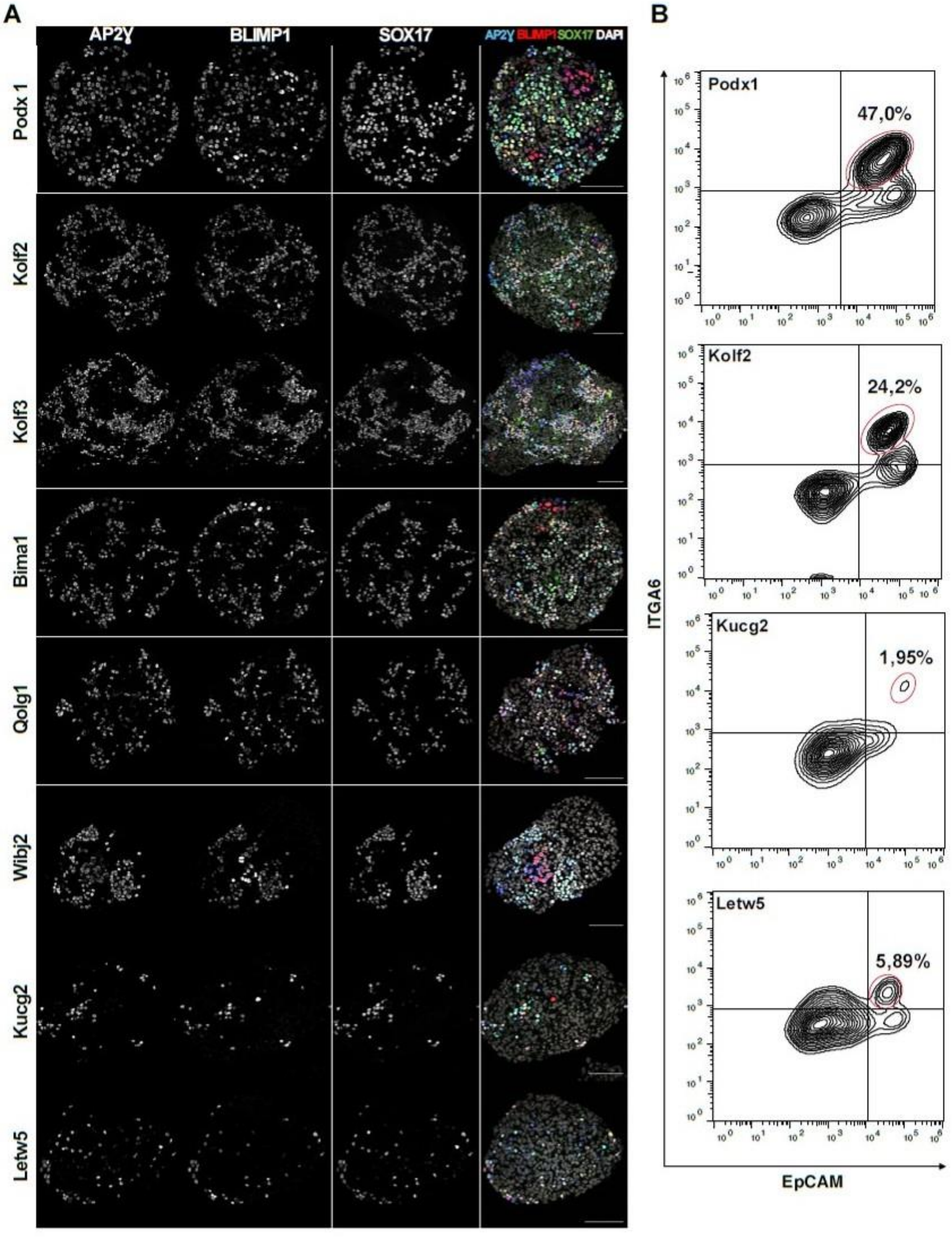

*Supplementary Figure 1.* ***Analysis of PGCLC differentiation efficiency.*** *A) Immunofluorescence images for AP2Ɣ, BLIMP1 and SOX17 of EBs at day 4 of differentiation from indicated hiPSC lines. AP2Ɣ+, BLIMP1+ and SOX17+ cells represent PGCLCs. B) FACS plots of day 4 EBs from indicated hiPSC lines stained for INTA6 and EpCAM. Population marked in red represents percentage of PGCLCs (INTA6+EpCAM+ cells) in EBs. Scale bars, 100µm.*

### Supplementary Figure 2

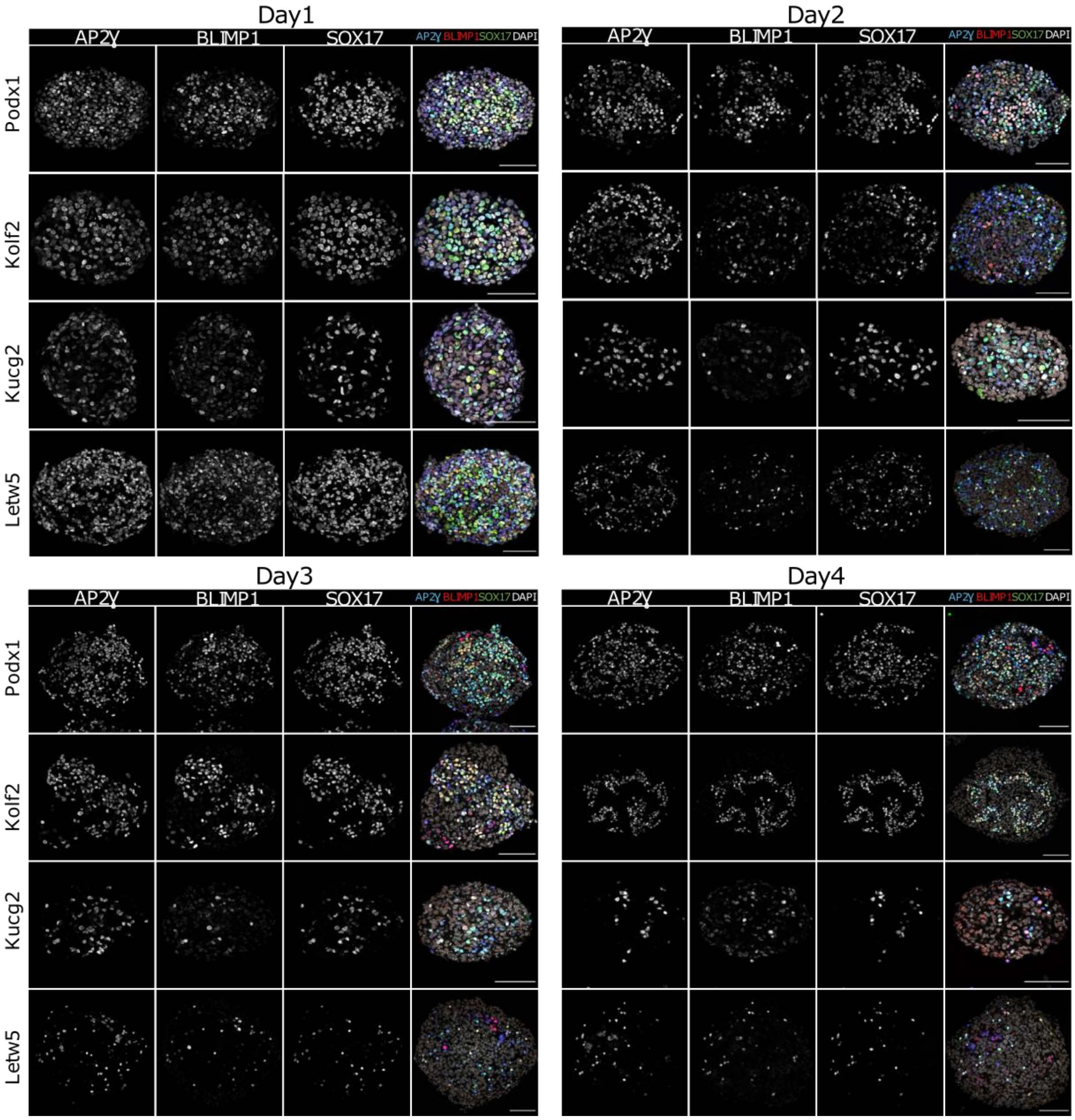

*Supplementary Figure 2.* ***Dynamics of PGCLC differentiation.*** *Immunofluorescence images for AP2Ɣ, BLIMP1 and SOX17 of day 1-4 EBs generated from indicated hiPSCs. Scale bars, 100 µm.*

### Supplementary Figure 3

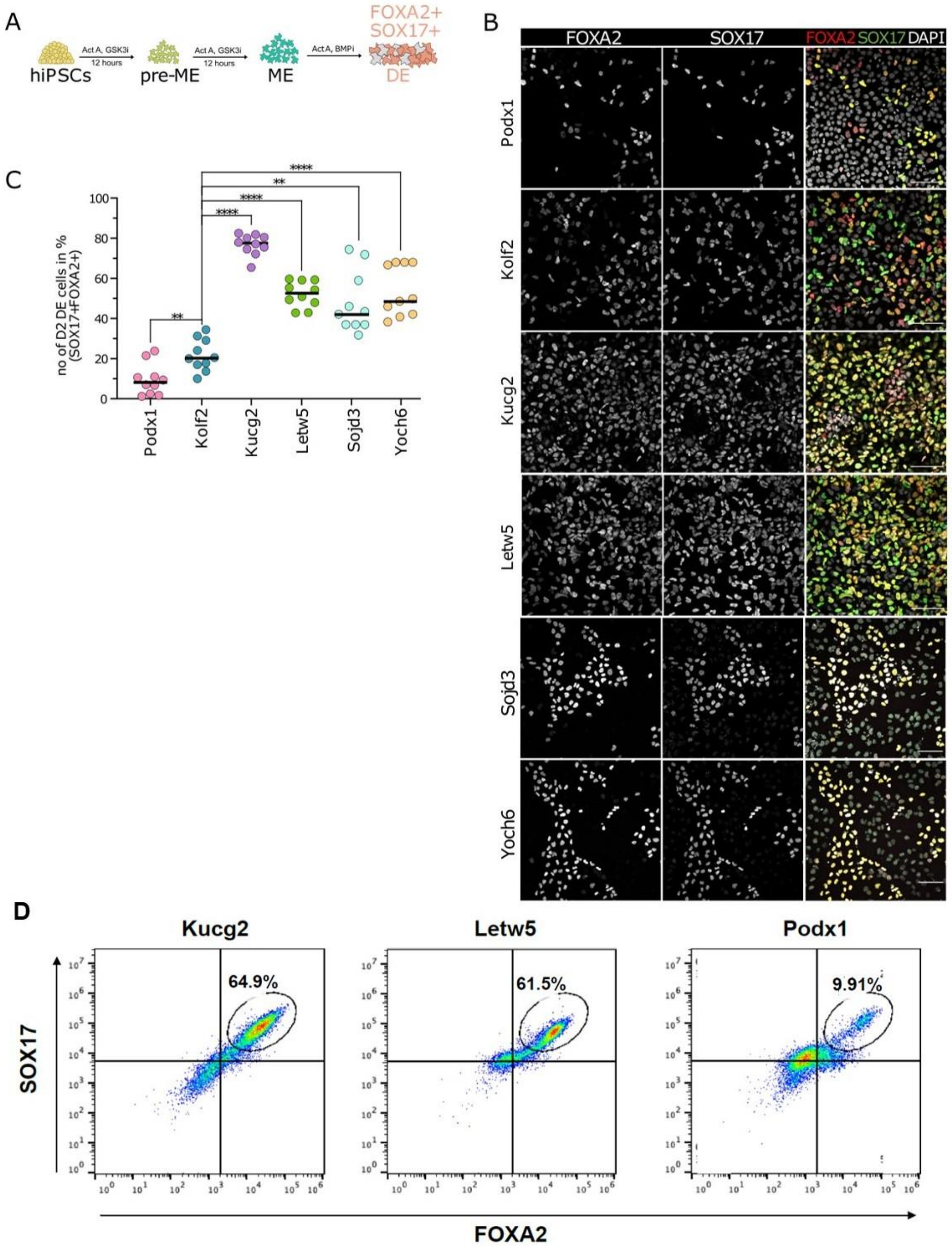

*Supplementary Figure 3.* ***Variation in DE differentiation efficiency****. A) Schematic representation of DE differentiation from hiPSCs. B) Immunofluorescence images for FOXA2 and SOX17 of DE cells after 48 hours of differentiation from indicated hiPSC lines. C) Quantification of immunofluorescence data after 2 days (D2) showing the percentage of FOXA2+ SOX17+ cells. Median values are represented by horizontal black lines. D) FACS analysis using SOX17 and FOXA2 staining of cell populations at day 3 of DE differentiation from Kucg2, Letw5 or Podx1 hiPSCs. SOX17/FOXA2-double positive cells (circled) indicate DE identity. Scale bars, 100 µm. Two-sided, unpaired t-test: **p-value <0.01; ****p-value <0.0001; ActA, Activin A; GSK3i, glycogen synthase kinase 3 inhibitor; pre-ME, precursors of mesendoderm; ME, mesendoderm; BMPi, bone morphogenetic protein inhibitor; DE, definitive endoderm.*

### Supplementary Figure 4

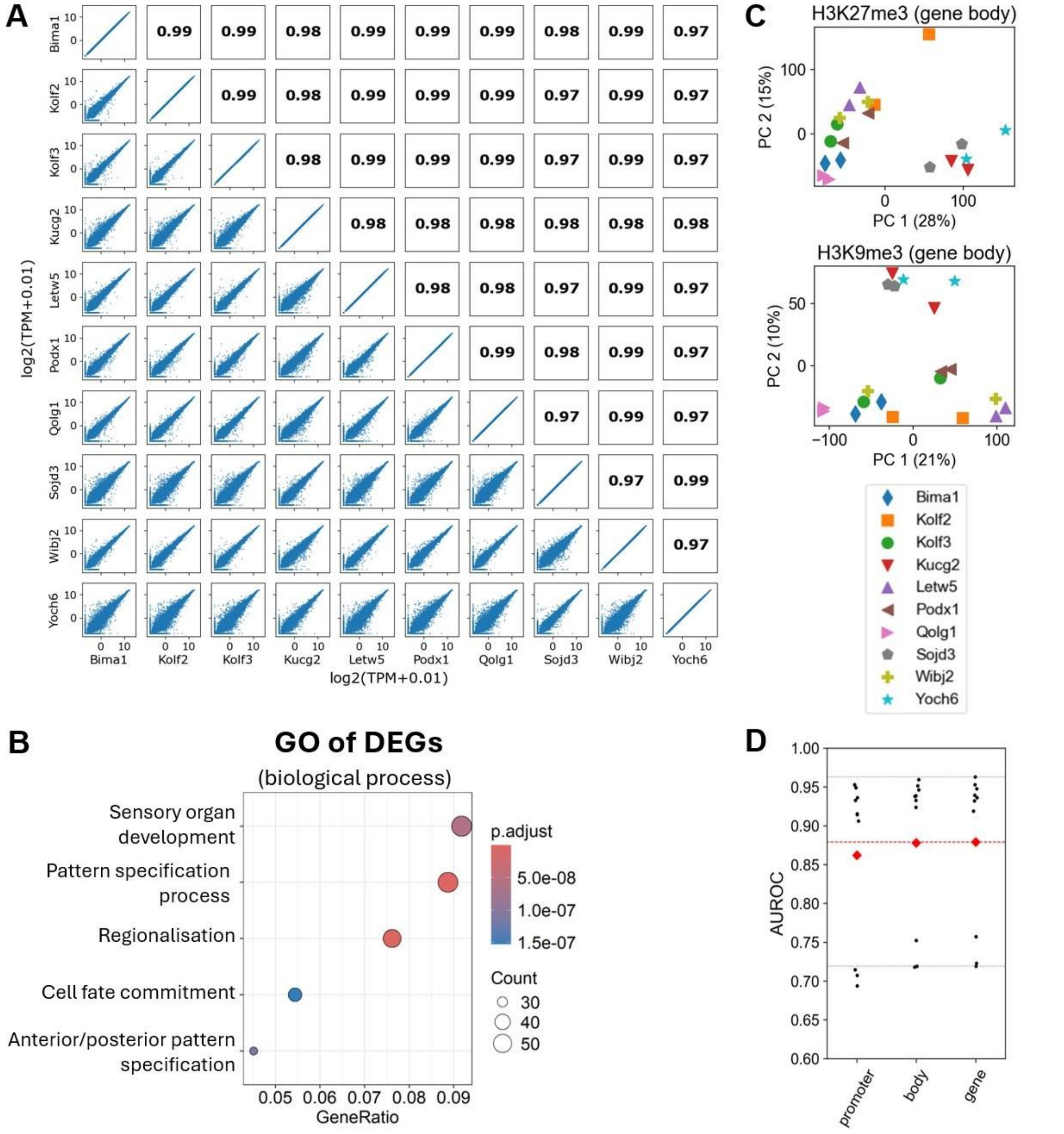

*Supplementary Figure* 4. ***Exploratory analysis of epigenomic data.*** *A) Scatterplots (lower half) and Pearson correlations (upper half) of the transcriptomes (autosomes only, in logarithmic scale) of the ten hiPSC lines. B) Gene ontology analysis of DEGs for biological processes. The fraction of genes (out of the total of 712) involved in a given molecular function is represented along the x-axis. C) PCAs of repressive histone marks in gene body annotations (from 1kbp downstream of gene start to gene end). The variance explained by each of the principal components is given in parenthesis, next to each axis. D) Area under the receiver operating characteristic (AUROC) for the SVM using different annotations for repressive marks (H3K9me3 and H3K27me3), with H3K4me3/accessibility using the standard ±1kb from the TSS promoter definition. Red diamonds correspond to the average AUROC, while the black dots represent the AUROC in individually tested cell lines. Dotted lines correspond to the average, maximum and minimum performance for the "gene" case (the one used in the manuscript).*

### Supplementary Figure 5

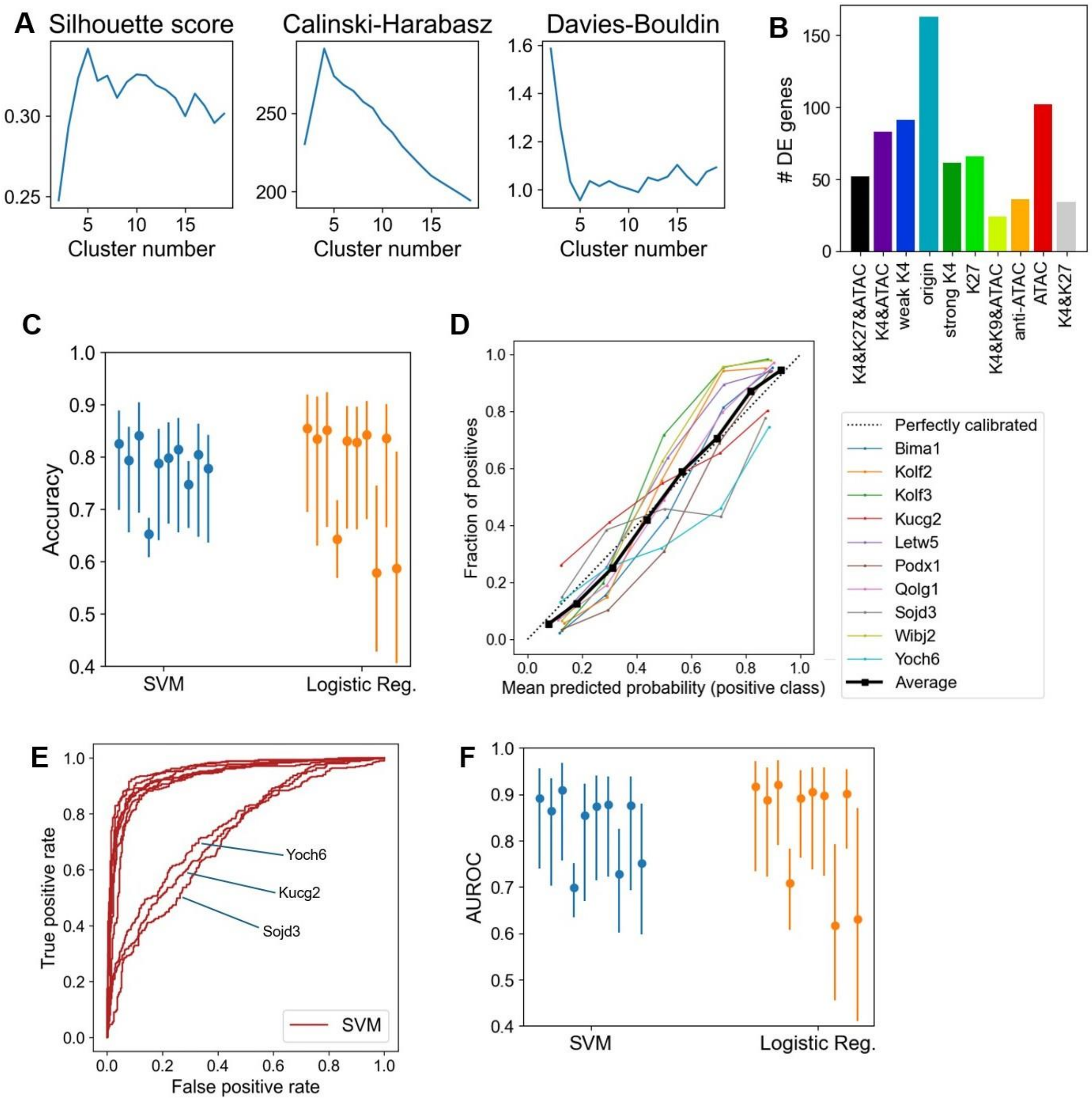

*Supplementary Figure 5.* ***Metrics of the SVM-based pipeline****. A) Goodness-of-clustering scores for the clustering into regulatory modes (for the first two, the higher the score the better the clustering; conversely for the rightmost score). B) Number of DEGs in each regulatory mode. C) Accuracy of the SVM (leave-one-out cross-validation, LOOCV). Filled circles show the mean value, and bars span along the 95% confidence intervals. Each of the bars corresponds to a line, which was withheld from the training set and whose behaviour we sought to predict (ordered alphabetically). For the alphabetical order of cell lines, see legend of panel D). Blue bars correspond to the accuracy using the SVM, while orange bars represent the accuracy of predictions using logistic regressions. D) Calibration plot for the SVM, binning the data into ten blocks. Dotted line represents a perfect calibration and "Average" corresponds to the overall calibration of the SVM, including all cell lines. E) ROC curve for the prediction of each of the ten hiPSC lines using the other nine lines (LOOCV), highlighting the three cell lines for which performance is lower. F) Distribution of AUROCs, for the SVM and logistic regression, shown as in panel C). Mean values and confidence intervals obtained from bootstrapping the training datasets $10^3$ times.*

### Supplementary Figure 6

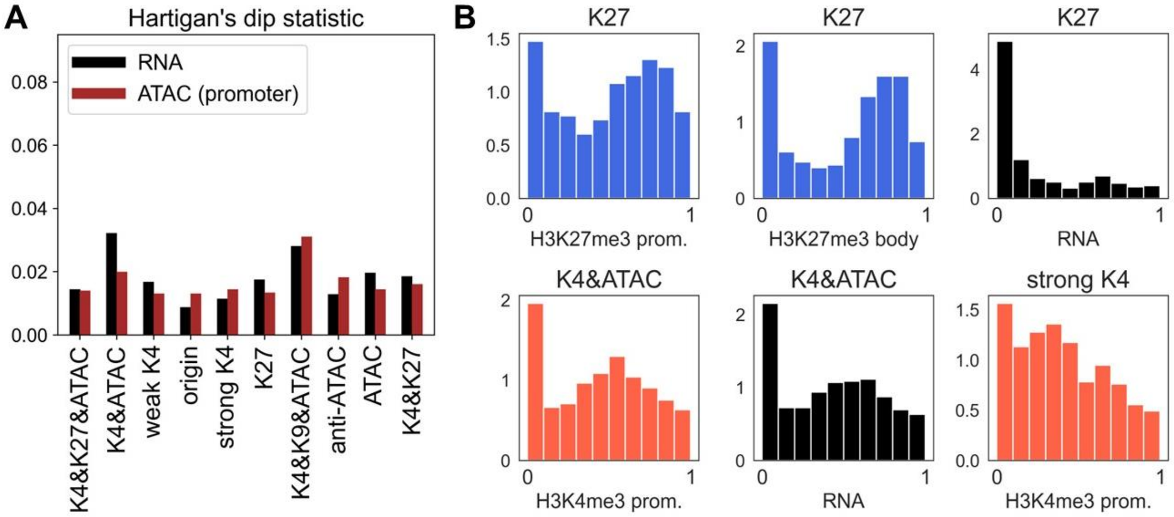

*Supplementary Figure 6.* ***Bimodality in epigenomic data.*** *A) Hartigan's Dip Statistic (HDS) for RNA (in TPMs, shown in black) and ATAC-seq signal at promoters (maroon), shown for each regulatory mode. B) Histograms of normalised values for a given variable (indicated on the x-axis) in a given regulatory mode (indicated in the title of each subpanel). One of them is strongly bimodal ('K27', H3K27me3 body), and two are not bimodal or very weakly bimodal ('K27', RNA and 'strong K4', H3K4me3 prom.). The other three represent intermediate cases.*

#### Supplementary Figure 7

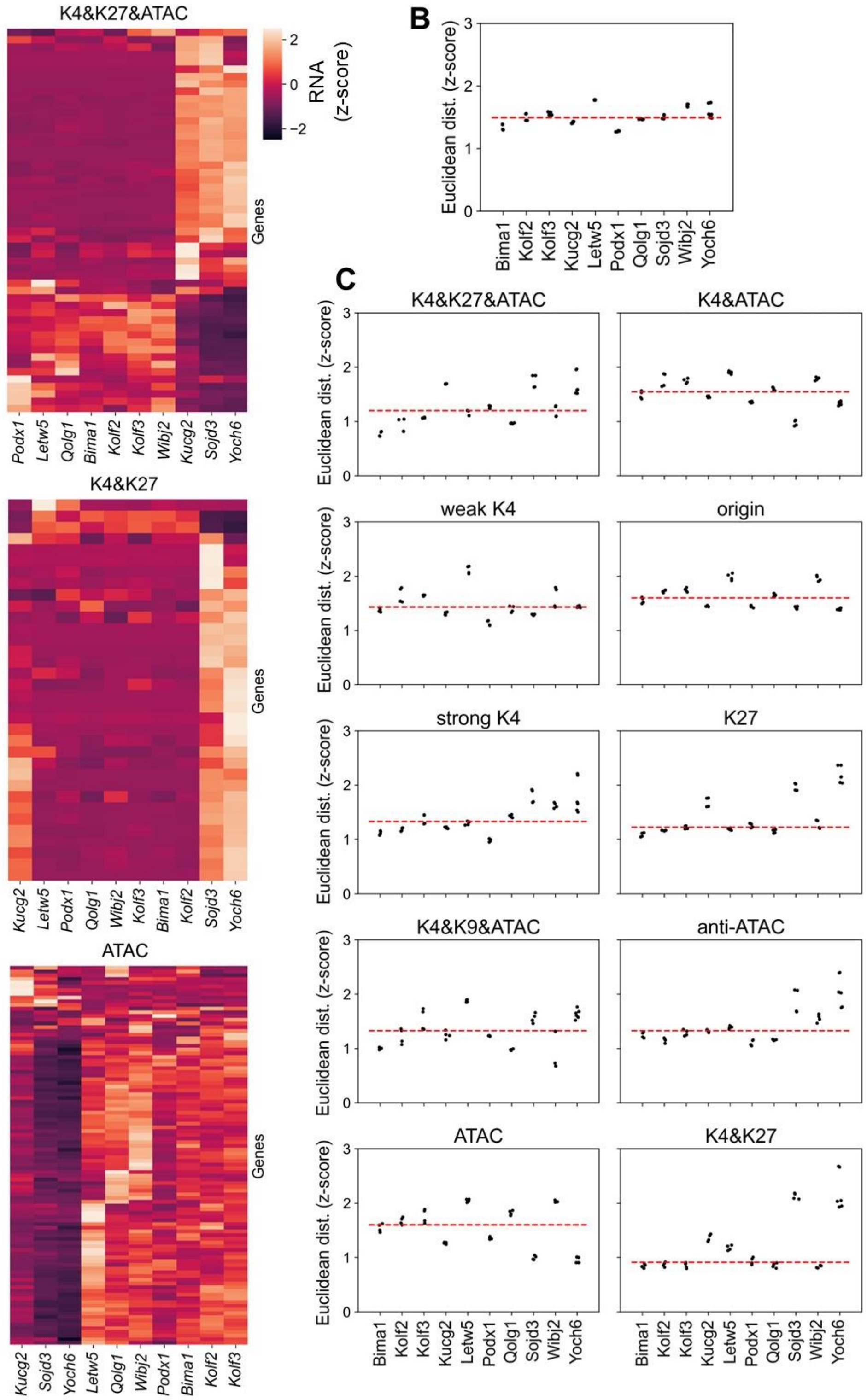

*Supplementary Figure 7.* ***Transcriptomic data dissected into regulatory modes****. A) z-score heatmaps of transcriptional outputs for three additional regulatory modes (shown as in Fig. 6A in the main text). B) Euclidean distances between the hiPSC lines and H9 hESCs using all DEGs. Red dashed line corresponds to the median distance between the hiPSC lines and the hESC line. C) Same as Fig. 6C in the main text, but for every regulatory mode.*

### Supplementary Figure 8

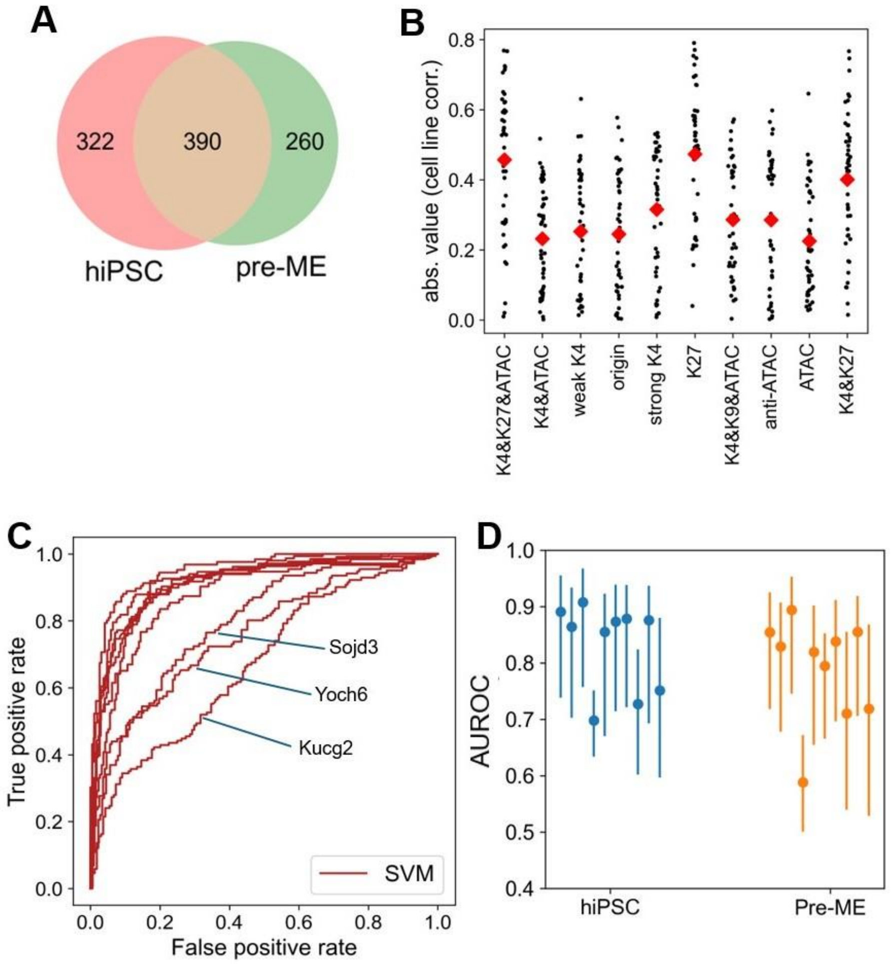

*Supplementary Figure 8.* ***Heterogeneities in pre-ME expression closely resemble those of hiPSCs.*** *A) Overlap in the DEGs among lines in hiPSC state and DEGs among lines in pre-ME state. B) Same as Fig. 6B in the main text but using pre-ME expression data. C) ROC curves for the performance of the SVM in predicting pre-ME expression in DEGs from chromatin variables in the hiPSC state. Same as Supplementary Fig. 5D, but for the prediction of pre-ME expression rather than hiPSC expression. D) Distribution of AUROCs, for the SVM to predict hiPSC expression or pre-ME expression from hiPSC chromatin variables. AUROC distributions shown as in Supplementary Fig. 5F. As expected, performance is reduced when using hiPSC features to predict pre-ME expression (with respect to predicting hiPSC expression), but it remains high overall.*
